# Bayesian network analysis incorporating genetic anchors complements conventional Mendelian randomization approaches for exploratory analysis of causal relationships in complex data

**DOI:** 10.1101/639864

**Authors:** Richard Howey, So-Youn Shin, Caroline Relton, George Davey Smith, Heather J. Cordell

**Author notes:** These authors contributed equally to this work (Richard Howey, So-Youn Shin). GNS Healthcare, 196 Broadway, Cambridge, MA 02139, USA.

## Abstract

Mendelian randomization (MR) implemented through instrumental variables analysis is an increasingly popular causal inference tool used in genetic epidemiology. But it can have limitations for evaluating simultaneous causal relationships in complex data sets that include, for example, multiple genetic predictors and multiple potential risk factors associated with the same genetic variant. Here we use real and simulated data to investigate Bayesian network analysis (BN) with the incorporation of directed arcs, representing genetic anchors, as an alternative approach. A Bayesian network describes the conditional dependencies/independencies of variables using a graphical model (a directed acyclic graph) with an accompanying joint probability. In real data, we found BN could be used to infer simultaneous causal relationships that confirmed the individual causal relationships suggested by bi-directional MR, while allowing for the existence of potential horizontal pleiotropy (that would violate MR assumptions). In simulated data, BN with two directional anchors (mimicking genetic instruments) had greater power for a fixed type 1 error than bi-directional MR, while BN with a single directional anchor performed better than or as well as bi-directional MR. Both BN and MR could be adversely affected by violations of their underlying assumptions (such as genetic confounding due to unmeasured horizontal pleiotropy). BN with no directional anchor generated inference that was no better than by chance, emphasizing the importance of directional anchors in BN (as in MR). Under highly pleiotropic simulated scenarios, BN outperformed both MR (and its recent extensions) and two recently-proposed alternative approaches: a multi-SNP mediation intersection-union test (SMUT) and a latent causal variable (LCV) test. We conclude that BN incorporating genetic anchors is a useful complementary method to conventional MR for exploring causal relationships in complex data sets such as those generated from modern “omics” technologies

**Author summary:** Mendelian randomization (MR) is a popular method for inferring causal relationships between variables (such as between an intermediate biological factor and a disease outcome). However, MR relies on a number of assumptions that may be hard to verify, and it is not ideally suited to comparing different underlying causal scenarios. Here we propose the use of an alternative approach, Bayesian network analysis (BN), as a complementary tool to conventional MR. We use real and simulated data to investigate the performance of MR, BN and several other recently-proposed methods, and find that BN performs as well as, or better than, the other methods, particularly under complex scenarios. We conclude that BN is a useful complementary approach to conventional MR for exploring causal relationships in complex data sets.

## Introduction

Causal inference methods offer an attractive avenue for understanding complex mechanisms in disease development and identifying ways to intervene upon them. An observed association between a risk factor and disease outcome does not necessarily imply causation, as it may arise via an alternative mechanism such as reverse causation or confounding [1]. A gold standard experimental approach for causal inference is to carry out a randomized controlled trial (RCT). By randomly allocating participants to intervention and control groups, an RCT can eliminate selection bias or confounding. However, it is an expensive and time-consuming process, and its result may imply a relatively short-term effect unless the trial is of long duration. Furthermore, intervention via an RCT is not always ethical, or (due to technical limitations) not feasible, for example when the potential risk factor involves DNA methylation or small metabolite variation.

Traditional non-experimental approaches for causal inference include discordant identical twin studies and longitudinal studies, which can be used to infer causal relationships under certain assumptions. Studies of identical twins are not subject to genetic confounding, and confounding by shared environmental factors is expected to be low (but reverse causation – which is unshared, and a major distorter of observational estimates — may bias the findings). In longitudinal studies with many repeated measures, methods such as g-computation can be applied [2, 3], but most longitudinal studies do not have the data measurements that allow the use of this approach.

### Mendelian Randomization

Mendelian randomization (MR) [4, 5] is an alternative non-experimental approach for causal inference applicable to a general population. In its simplest form it utilizes a genetic variant whose robust association with a risk factor provides a directional causal anchor. The approach is based on the fact that there is only one fixed direction of causation between the genetic variant and the outcome. Use of the genetic variant (which is allocated at gamete formation during conception) has some analogies to the randomization procedure in an RCT. Hence, causal inference is made from the difference in the outcome seen between people with different genetic variants. MR was not originally introduced as an instrumental variable (IV) approach [4, 6], and genetic anchors can contribute to causal inference in some settings in which formal IV analysis cannot be applied. However, the main contemporary implementation of MR is in analyses in which genetic variants, usually single nucleotide polymorphisms (SNPs), are considered to operate as IVs, provided certain conditions are met [7, 8]. Henceforth in this paper we use the abbreviation “MR” to refer to the now-conventional approach of applying IV analytical strategies to evaluate the evidence for causal relationships between genetically instrumented risk factors and phenotypic outcomes.

MR has been widely applied to evaluate the causal role of traditional risk factors in disease, such as HDL and LDL cholesterol in cardiovascular disease [9, 10]. It has also been applied to identify innovative drug targets in early stage drug development or to discover novel risk factors at the molecular level by scanning “omics” data systemically [11–13]. An advantage of MR is that individual-level data are not necessarily required; inference can be performed on the basis of summary statistics measuring the relationship between the genetic instrument(s) and the risk factor, and the relationship between the genetic instrument(s) and the outcome [14]. This means that the summary statistics required to perform MR analysis can be derived from different studies, in an approach termed two-sample MR [15–17].

Nevertheless, MR has limitations. MR works only if there is a genetic variant robustly associated with the risk factor. It has relatively low statistical power and thus requires a large sample size. MR also has drawbacks in analysis of large-scale “omics” data. Such data often include a number of measured traits that are highly correlated with each other, and some of these may be associated with the same genetic variant(s) and the same outcome. If MR were naively applied for each of these correlated traits, this could violate the MR assumption that the genetic variant used as an instrument influences the outcome only via the risk factor tested.

To address this issue, several approaches that attempt to either detect or allow for pleiotropy in the context of MR, or to investigate more complex networks of relationships between variables, have been proposed [18–26]. There has also been considerable interest in using MR in a “bi-directional”’ or “reciprocal” fashion to determine the direction of causation between two variables, say X and Y [15, 27]. In most of these approaches, an underlying hypothesized graphical structure representing the relationships between variables must be assumed (rather than being learned from the data). However a recently-proposed addition to bi-directional MR, known as MR Steiger [28], moves a step further by first carrying out an initial determination of whether a genetic variable G is most suitable as an IV for variable X or Y, prior to conducting standard a MR analysis between them based on the determined relationship. This use of Steiger filtering in the context of bi-directional MR is an important component that improves correct directional identification. Another recent method [29] addresses a similar goal through use of a latent causal variable (LCV) model to infer, for all pairs of traits of interest, the extent to which part or all of the genetic component of one trait is causal for another, suggesting (although not formally demonstrating) that one trait may itself be causal on the other.

### Bayesian Network Analysis

Bayesian networks can be used as the basis for an alternative set of non-experimental, statistical techniques for causal inference. First formalized and developed by Pearl [30, 31], Bayesian networks have now become widely applied in the social and natural sciences. Briefly, a Bayesian network describes the conditional dependencies of variables using a graphical model known as a directed acyclic graph (DAG) and an accompanying joint probability [32]. In a DAG, the variables and their conditional relationships are represented as nodes and directional arcs or edges (arrows), respectively. The joint probability is decomposed as a product of local probabilities where the local probability of each variable is explained by its conditional dependencies on its immediate neighbours [33]. The local probability distribution can take any form, but usually a multinomial distribution is used for discrete variables and a multivariate normal distribution is used for continuous variables.

Although Bayesian networks are generally considered to encapsulate conditional dependencies (and independencies) between variables rather than necessarily implying causal relationships between them – resulting in a distinction between (probabilistic) “Bayesian networks” and “causal Bayesian networks” [34] – a causal interpretation can be ascribed to a Bayesian network under three assumptions [32]: 1) the causal Markov assumption, 2) the causal faithfulness assumption, and 3) the causal sufficiency assumption. The causal Markov assumption states that a variable is independent of all other variables, except for its effect or descendent (“child”/“grandchild” etc.) variables, conditional on its direct causal (or “parent”) variables [32, 33, 35]. The causal faithfulness assumption states that the network structure and the causal Markov relations assumed represents all (and the only existing) conditional independence relationships among variables [32, 36]. The causal sufficiency assumption corresponds to asserting that there are no external variables which are causes of two or more variables within the model, implying that all causes of the variables are included in the data and there are no unobserved confounding variables [32, 36, 37]. A further (sometimes unappreciated) assumption is that of no measurement error i.e. the variables are measured without any errors [36]. These assumptions are essential for causal inference, and are quite commonly assumed in other causal inference methods, but they are generally impossible to validate (and, indeed, may be considered unlikely to hold completely, raising the question of sensitivity to their violation). In the MR literature a large (and growing) set of sensitivity analyses allow relaxation of some of the assumptions required for identification [38].

In Bayesian network analysis (BN), we try to infer the most plausible DAG (or a set of plausible DAGs), and thus potential causal relationships between variables, given data (measurements of the relevant variables). In this paper we use the abbreviation “BN” to refer (in principle) to a whole suite of different analysis approaches (employing different methods/algorithms) that use Bayesian networks for the purpose of inferring the most plausible DAG(s), although in our simulations and real data analyses we focus primarily on one specific (score-based) algorithm (see Methods). Various algorithms have been developed to infer the DAG that best fits the data [39]. Constraint-based methods start with a fully connected graph and carry out a series of marginal and conditional independence tests to decide which edges to remove. These methods can be considered non-parametric, as they focus on testing conditional independence rather than requiring specification of a parametric likelihood. Score-based methods require a likelihood and hence are parametric (unless all nodes are discrete); they move around through graph space in order to determine the most plausible DAG i.e. the DAG that has the best score (highest or lowest, depending on how the score function is defined), or the highest posterior probability, out of all possible DAGs. As the number of variables in the data set increases, the number of all possible DAGs increases and the enumeration of all possible DAGs becomes infeasible [40]. Thus, in many cases, the most likely DAG is estimated using a model search algorithm or a model averaging algorithm. As the DAG structure is learned/estimated, the parameters of the probability distributions are also learned/estimated from the data using a parameter search algorithm such as maximum likelihood estimation or Bayesian estimation. Note that the term “Bayesian” within BN refers to the fact that certain calculations rely heavily on Bayes’ theorem, rather than implying the use of a Bayesian – as opposed to a frequentist – paradigm.

Intuitively, one would expect the search algorithms used in BN to perform better when directional anchors are available. Directional anchors prevent edges from coming towards certain nodes, which reduces the number of all possible DAGs dramatically and improves the model search process by distinguishing one possible DAG out of a statistically equivalent class of DAGs [36]. In analysis of “omics” data, genetic variants are natural instruments that can be used to define directional anchors.

BN has the ability to accommodate large complex data relatively flexibly. This feature is particularly useful when the study aims to address simultaneous causal relationships in “omics”-scale data sets, for example in studies of gene expression [41] or metabolites [42]. Recent network-building methods have been developed that allow the analysis of potentially hundreds of variables, including both discrete and continuous data types, taking advantage of the ability of genetic variables to operate as causal anchors to help orient the direction of relationships between non-genetic variables [43–47]. These BN approaches could be considered complementary to the MR-based approaches [28] that enable the construction of such networks, as they use very different algorithms, are more restrictive in terms of requiring individual level input data, and produce different outputs, albeit in order to achieve similar goals. Nevertheless, BN has known limitations. The model search process for score-based algorithms, particularly with large numbers of variables, requires massive computational power and often elimination or pre-filtering of variables is required. The causal relationships implied by each tested model are only strictly valid in the (somewhat implausible) “no measurement error, no unmeasured confounding” situation (though this assumption may sometimes be defended by appealing to “prior background knowledge”); any violation of the assumed relationships will affect the calculation of the penalized log likelihood (or other criterion) used as the basis of the network scores. In particular, in common with several causal inference methods, results from BN may be biased in the presence of hidden confounding factors. This is due to violation of the causal sufficiency assumption.

## Results

We applied MR and BN approaches to both real and simulated data, in order to investigate the properties and performance of the different approaches. Our simulations included scenarios where the assumptions for valid causal inference (using MR and/or BN) were violated, as we were interested in investigating the extent to which this would result in erroneous or misleading inferences. We also applied two recently-proposed methods, LCV [29] and SMUT [48], along with BN, MR and a recent MR extension [25], to data generated under a more complex simulation scenario involving extreme pleiotropy.

### Illustrative Example: fatty acids and BMI

As an initial motivating example, we investigated possible causal relationships between fatty acid metabolites and body mass index (BMI) using the TwinsUK study data [49]. We applied both MR and BN to these data, and compared the causal inferences obtained. We note that this example is intended as a (relatively straightforward) illustration of analysing data using both MR and BN approaches, rather than making any strong claims for the validity of the instruments (and thus for the robustness of the inferences obtained) in this particular case.

The metabolites considered were the omega-3 fatty acids eicosapentaenoate (EPA) and dihomo-linolenate (DGLA). For testing whether a causal relationship existed between these metabolites and BMI, genetic IVs for EPA and DGLA were chosen based on knowledge gained from prior investigation of this data set (along with an additional German cohort) [50]; we note that re-use of (some of) the same data used to identify the instruments can, in theory, run the risk of over-fitting. Based on these previous results, the SNP rs174556 in *FADS1-2-3* was used as an IV for EPA, while the SNPs rs968567 in *FADS1-2-3* and rs6498540 in *PDXDC1* were used as IVs for DGLA.

rs174556 and rs968567 are in linkage disequilibrium (LD) i.e. correlated with an *r*^2^ value of ≈ 0.52. It is conceivable that rs174556 is actually a causal variant for DGLA, and so could have an effect on BMI operating in parallel through both EPA and DGLA. This would violate one of the three assumptions required for the genetic variant to be used as an instrumental variable (IV) for EPA, namely the no-horizontal pleiotropy assumption that the IV has no effect on the outcome besides the effect mediated through the risk factor (EPA).

MR, based on the individual level data (rather than based on summary statistics via two-sample MR), was used to test for a causal relationship between each metabolite and BMI. The rationale for using individual level data (rather than performing the asymptotically equivalent two-sample MR analysis) was to allow comparison with BN which (at least in its current implementations) requires access to individual level data. A causal relationship from BMI to each metabolite was also tested using MR with an instrumental variable for BMI given by a BMI allele score formed (on the basis of prior knowledge [51, 52]) from 39 BMI-associated SNPs. Again, individual level data (and resulting individual level BMI allele scores) were used, although the weighting of the SNPs to construct the allele score variable could be considered to incorporate external information as “prior knowledge”, being informed by previous results from larger studies [51, 52].

Table 1 shows the results of applying Mendelian randomization to the TwinsUK data set. Both metabolites, EPA and DGLA, were inferred to have a causal relationship with BMI at the 0.05 significance level (*p*-values 0.0498 and 0.0327 respectively). Conversely, reverse causation (with BMI causing the metabolite levels) did not show such compelling *p*-values (0.6668 and 0.7082 respectively). Mendelian randomization between the metabolites provided somewhat conflicting results, with both directions achieving *p*-values *<* 0.05, but overall there seemed to be slightly stronger support for a causal effect from DGLA to EPA. Since rs174556 is used as an instrument for EPA, while rs968567 and rs6498540 are used simultaneously as instruments for DGLA, the LD between rs174556 and rs968567 violates the exclusion restriction (that an instrument only influences the outcome through the exposure) in both directions, complicating the interpretation of the results from MR, and thus perhaps motivating use of an alternative method such as BN, which avoids the problem of having an additional (unaccounted for) path between a genetic instrument and the outcome by simultaneously modelling the effects of all measured instruments and exposures.

**Table 1.**
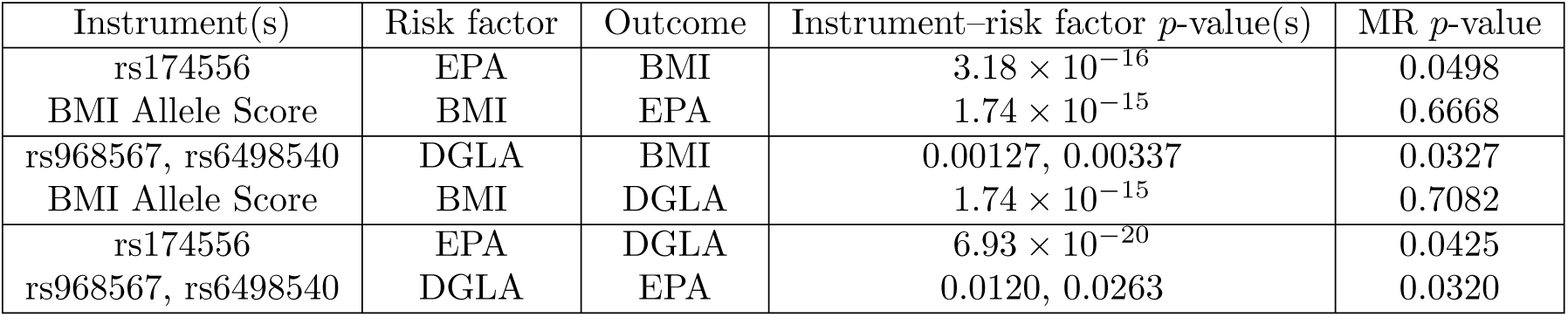
Mendelian randomization results for TwinsUK data. The Instrument–risk factor *p*-value(s) are from the regression of the risk factor on the instrument(s), and the MR *p*-value is from the regression using the predicted value of the risk factor as an explanatory variable for the outcome variable, with the uncertainty in the predicted values from the first-stage regression taken into account through use of the R function ivreg().

Fig 1A shows the average network from BN when all variables are included, with the directional constraint that arrows are constrained to come out from (rather than go into) any nodes corresponding to genetic variables. The thickness of the edges indicates their strength or probability of existence (i.e. frequency of edge presence in all replicates), providing a visual representation of the relative support for the possible causal effects. The red numbers indicate the probability of existence of the edge, and the numbers in brackets indicate the probability of the edge operating in direction shown, given that it exists. The edges for which both numbers are provided are those in which we are most interested, namely those that represent relationships between BMI and the metabolites. The other edges are annotated with only one number, their probability of existence, as they were already constrained such that edges from the SNPs and the BMI score variables could only go in one direction (outwards from the variable towards a child node), consistent with the notion of these variables acting as genetic instruments or anchor variables. The average network shows strong evidence (overall probabilities of 0.89 = 0.96 × 0.93, and 0.86 = 0.99 × 0.87, respectively) of DGLA and EPA being causal on BMI.

**Fig 1.**
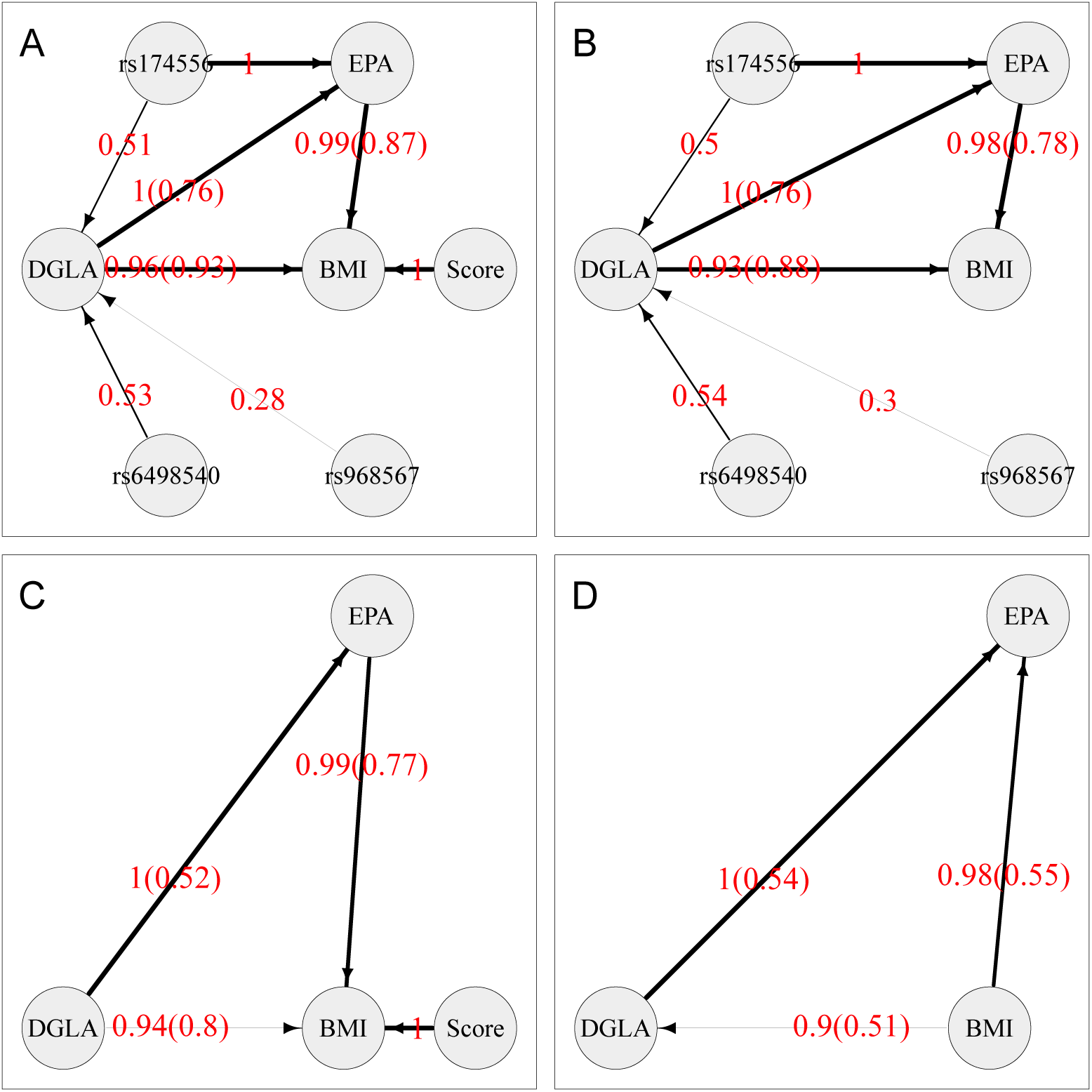
Average Bayesian networks for the TwinsUK data using either (A) all available variables or (B, C, D) a subset of variables, as shown. The red numbers indicate the probability of existence of the edge, and the numbers in brackets indicate the probability of the edge operating in direction shown, given that it exists. The thickness of an edge indicates its strength (probability of existence).

With the removal of the BMI score variable (Fig 1B), the probabilities are slightly decreased to 0.82 = 0.93 × 0.88 and 0.76 = 0.98 × 0.78, still supporting the direction of relationships between the metabolites and BMI when one instrument (the BMI score) is removed. Similarly, when only the BMI score anchor variable is present (Fig 1C), the relationships between the metabolites and BMI are still supported, although with reduced probabilities of 0.75 = 0.94 × 0.80 and 0.76 = 0.99 × 0.77. However, when all instruments are removed (i.e. only variables DGLA, EPA and BMI are included in the analysis) (Fig 1D), the direction-of-edge probabilities between the three variables are all close to 0.5, illustrating the fact that, in the absence of any instrumental variables to anchor the network, all networks connecting all three variables are statistically equivalent, thus the likelihoods are equal and no directional preference can be determined.

Similarly to MR, BN also suggests a causal relationship from DGLA to EPA, with the estimated probability of the direction decreasing as the number of anchor variables is reduced.

The fits of the four models shown in Fig 1 A–D are not directly comparable, as they contain (and thus model data at) different numbers of variables. However the fits of models that do or do not contain arrows between the genetic variables and the non-genetic variables can be examined by fitting networks equivalent to that shown in Fig 1A (so modelling data at all measured variables), but with certain arrows “blacklisted” to not be allowed to exist. The average network score based on the Bayesian information criterion (BIC) when all variables are included and both the SNPs and the BMI allele score are allowed to have children (equivalent to Fig 1A) is −33500.99. The average network score when all variables are included and SNPs can have children but the BMI allele score is constrained to have no children (conceptually similar to the model in Fig 1B) is −33528.85. The average network score when all variables are included and the BMI allele score can have children but the SNPs are constrained to have no children (conceptually similar to the model in Fig 1C) is −33546.05. The average network score when all variables are included and SNPs and the BMI allele score are both constrained to have no children (conceptually similar to the model in Fig 1D) is −33573.91. These average BICs illustrate the considerably better fit obtained when all anchor variables are allowed to influence the values of the other variables in the model.

Overall, these results support the inference seen with this data set using MR. They also illustrate the advantage in BN of being able to easily include simultaneously anchor variables for both BMI and the metabolites – although removing one or other anchor still produced broadly similar inference concerning the direction of the relationships between the metabolites and BMI, we found the support for these relationships (as measured by the estimated probabilities of the directions of the relevant edges, see Figs 1B and 1C) was lowered.

In this example, we did not use BN to directly model LD between SNPs. It would, in fact, be possible to construct BN models that incorporate LD between variants by allowing arrows to connect genetic variables, with the practical constraints that arrows are only allowed between SNPs that lie relatively close together on the genome, and the directions of the arrows are constrained to go only in one direction (e.g. from smaller to larger base-pair positions). Whether this would offer any practical advantage in terms of elucidating the relationships between the non-genetic factors, or between genetic and non-genetic factors, would be an interesting topic for further investigation.

A different consequence of LD is the fact that none of the SNPs being used as instruments for EPA or DGLA are necessarily causal for the fatty acids (or for BMI); they may simply be surrogates for (i.e. correlated with) the true unmeasured causal variants. This would not be expected to invalidate the analysis for either MR or BN, but would result in weaker relationships between the measured SNPs and the fatty acids, and thus lower power for detection of network relationships. It is even conceivable that there is a single causal SNP affecting both EPA and DGLA, which is tagged to different extents by the three measured SNPs – although this seems unlikely in view of the fact that, in Fig 1A, rs968567 and rs6498540 are chosen as instruments for DGLA but not for EPA, while rs174556 is chosen as an instrument for both DGLA and EPA. (If there were, in fact, a single causal SNP affecting both EPA and DGLA, one would expect the same surrogate instruments, namely those most strongly correlated with the causal SNP, to be chosen for both EPA and DGLA). In principle, one could fit a graph with an additional latent node for an unmeasured causal SNP in order to explore this scenario. Although not explored here, causal discovery methods such as the Fast Causal Inference (FCI) algorithm, that allow latent (unmeasured) nodes resulting in non-directed, bi-directed or partially-oriented edges between measured variables, do exist [33], and would be an interesting topic for further investigation.

### Simulation Study 1

#### MR and BN powers and type I errors

We initially used three relatively simple simulation models (Fig 2) to investigate the powers and type I errors of MR, MR Steiger and BN for testing the relationship X to Y (or Y to X), with assumed effect size *β_XY_* or *β_Y X_* = 0.5 (or 0). By definition, MR with G used as an IV for X tests the relationship X to Y (and MR with Z used as an IV for Y tests the relationship Y to X), while MR Steiger chooses the most appropriate instrument (G or Z) to use as an instrument when testing the relationship X to Y (or Y to X). We implemented MR and MR Steiger using instrumental variable regression, which takes into account the uncertainty of the predicted values in the first-stage regression (see Methods). We also compared the results with those obtained (denoted MR’ and St’) using two-stage least squares regression without accounting for the uncertainty of the predicted values in the first-stage regression, which provides equivalent inference (identical p-values) to that obtained by simply regressing the outcome variable on the chosen IV. For BN, the existence (marginal posterior probability) of an edge going from X to Y (or Y to X) is determined by counting the proportion of times that the relevant edge occurs amongst 1000 bootstrap networks (see Methods).

**Fig 2.**
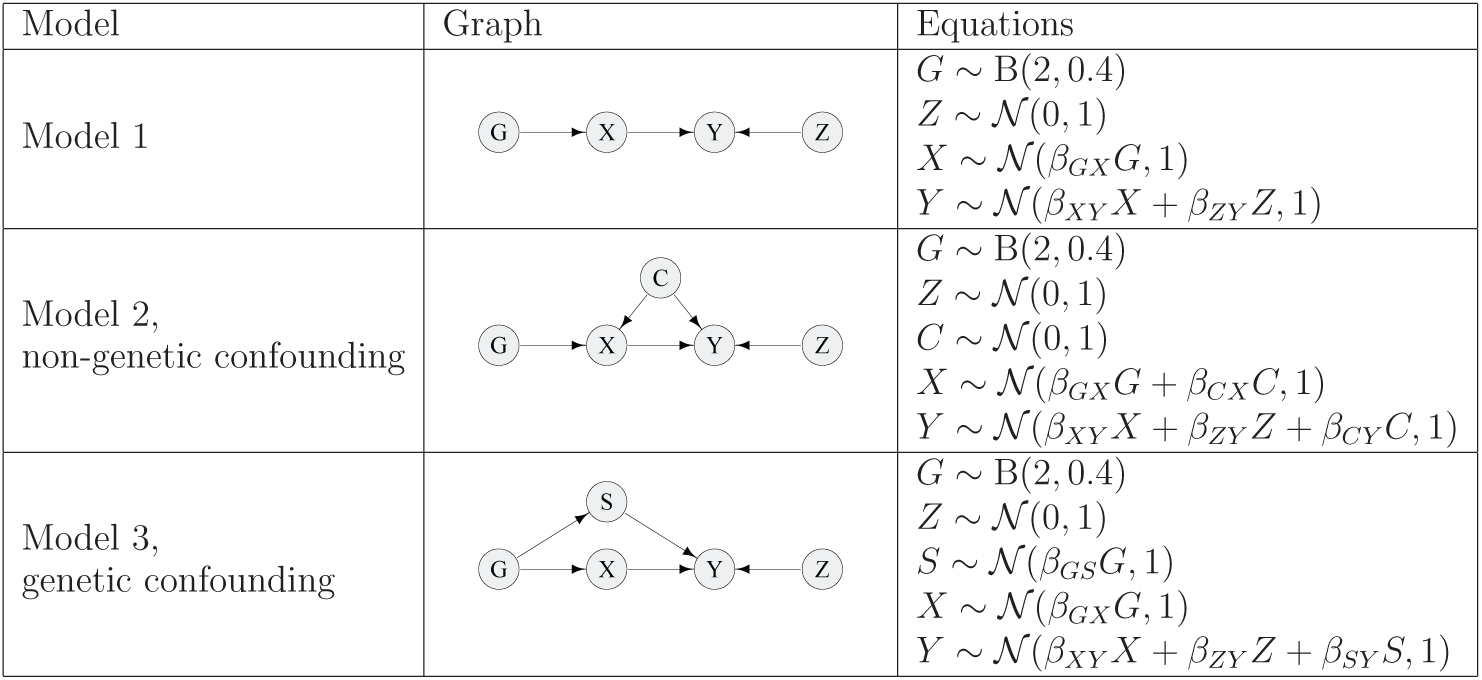
Simulation models used in Simulation Study 1 of quantitative trait data. Data were simulated for two continuous variables (X and Y), together with a genetic instrument G (coded as 0, 1, 2) and a continuous instrumental variable Z. Parameter values for models involving weak confounding were chosen as *β_GX_* = 0.1, *β_ZY_* = 0.075 and *β_CX_* = *β_CY_* = *β_GS_* = *β_SY_* = 0.25. For models involving strong confounding, the parameter values were the same except that *β_CX_* = *β_CY_* = *β_GS_* = *β_SY_* = 0.5 i.e. the parameters controlling the confounding effects were doubled. The parameter *β_XY_* was varied using values of 0.0, 0.1, 0.2, 0.3, 0.4 and 0.5. For the calculations where false positives were counted as detections of an arrow between X and Y in the wrong direction, the direction of causality was reversed between *X* and *Y*, such that for model 1 the equations become *X ∼* N (*β_Y X_Y* + *β_GX_G*, 1) and *Y ∼* N (*β_ZY_ Z*, 1), with *β_YX_* varied using values of 0.1, 0.3 and 0.5 (and similarly for models 2 and 3).

The results shown in the left hand plots (model 1) of Fig 3 and 4 involve no confounding, the results in the middle and right hand plots (models 2 and 3) of Fig 3 involve weak confounding, while those in the middle and right hand plots of Fig 4 involve strong confounding. More detailed results under weak confounding for testing either X to Y or Y to X (given an effect *β_XY_*, with different effect sizes) using MR and MR Steiger are shown in Figs S1-S3, with a comparison between MR, MR Steiger and BN shown in Figs S4-S6 (for BN implemented via the **bnlearn** algorithm [39]) and Figs S7-S9 (for BN implemented via the **deal** algorithm [53]). Comparison of Figs S4-S6 with Figs S7-S9 shows the power of the **deal** algorithm to be consistently lower than that of the **bnlearn** algorithm (with no compensating advantage in terms of better type 1 error), and for this reason we discard the **deal** algorithm from any further consideration.

**Fig 3.**
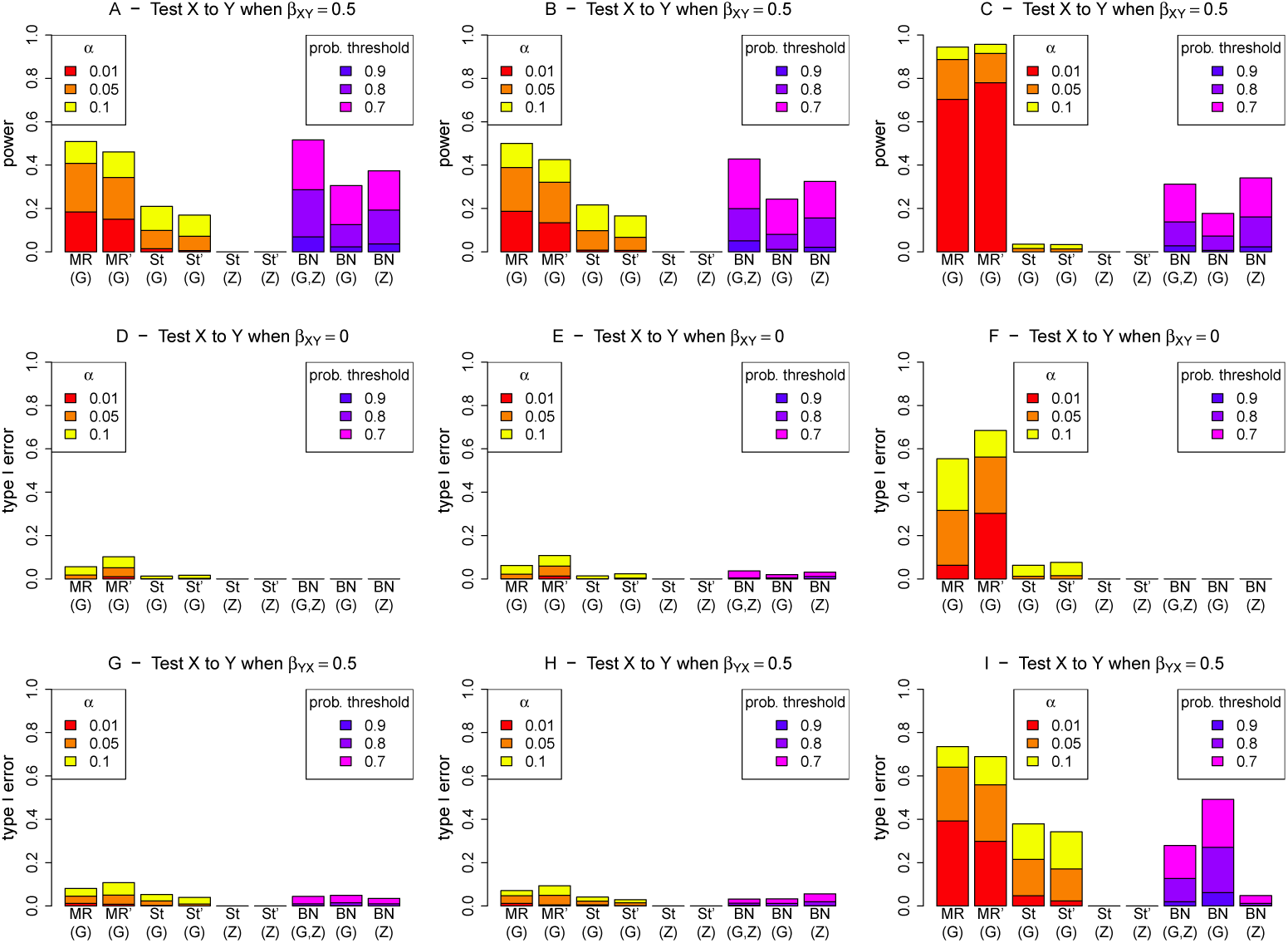
Performance (power and type I error) of different methods for detecting an edge from X to Y, under different generating scenarios that include weak confounding. MR and St denote MR and MR Steiger respectively, performed using instrumental variable regression which takes into account the uncertainty of the predicted values in the first-stage regression to calculate the MR *p*-values. MR’ and St’ denote MR and MR Steiger respectively, performed using two-stage least squares regression without accounting for the uncertainty of the predicted values in the first-stage regression. Left hand plots (A, D, G) are generated under model 1 (no confounding), middle plots (B, E, H) are generated under model 2 (non-genetic confounding), and right hand plots (C, F, I) are generated under model 3 (genetic confounding).

**Fig 4.**
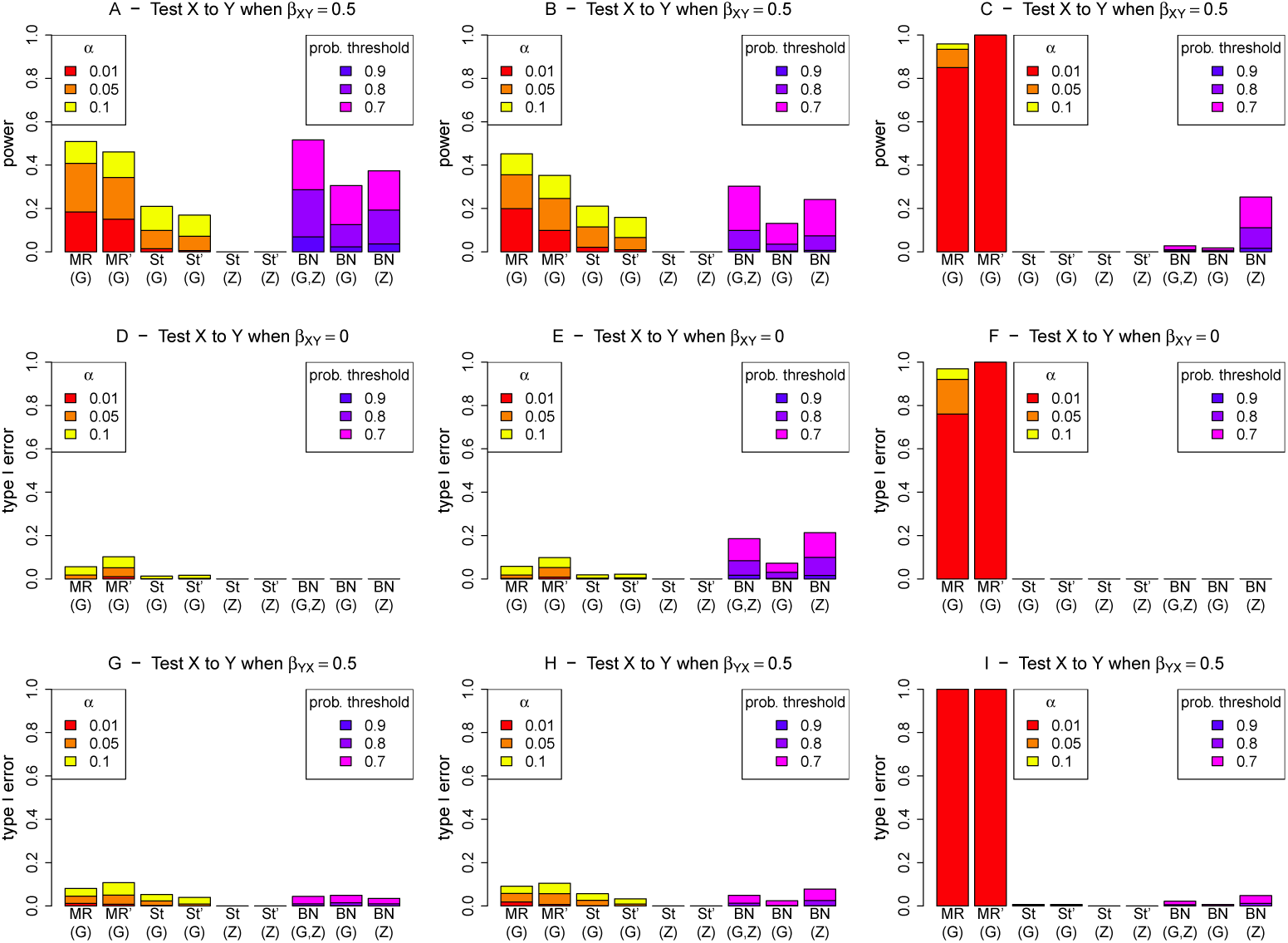
Performance (power and type I error) of different methods for detecting an edge from X to Y, under different generating scenarios that include strong confounding. MR and St denote MR and MR Steiger respectively, performed using instrumental variable regression which takes into account the uncertainty of the predicted values in the first-stage regression to calculate the MR *p*-values. MR’ and St’ denote MR and MR Steiger respectively, performed using two-stage least squares regression without accounting for the uncertainty of the predicted values in the first-stage regression. Left hand plots (A, D, G) are generated under model 1 (no confounding), middle plots (B, E, H) are generated under model 2 (non-genetic confounding), and right hand plots (C, F, I) are generated under model 3 (genetic confounding).

Direct comparison between MR/MR Steiger and BN is complicated as the most natural processes for determining if X is causal on Y (or for examining the extent of the evidence for X being causal on Y) are different for the three methods. For MR/MR Steiger, we make inference based on a type I error (*p*-value) threshold given by *α* (the probability of a false positive). Although we concur with the opinion [54] that, in real data analysis, use of absolute *p*-value thresholds should be avoided, we do not in fact propose that any particular threshold should be considered as “correct”; here we instead use the thresholds as heuristics and examine the performance of the methods (in terms of true and false detections of relationships) as the thresholds are varied. For BN, we make inference based on an estimated posterior probability that X is causal on Y, and again examine performance of the method as the threshold for declaring detection is varied. Given the lack of direct comparability between the methods, we use different thresholds (and different colours) for MR/MR Steiger and BN in the plots shown in Fig 3-4 and Figs S1-S9: for MR/MR Steiger we use *α* values of 0.01, 0.05 and 0.1, while for BN we use probability thresholds of 0.7, 0.8 and 0.9. The resulting powers and type I errors are therefore not directly comparable, but they do give some indication of how the methods perform using thresholds that might be considered reasonable choices in practice.

For MR performed using instrumental variable regression, which takes into account the uncertainty of the predicted values in the first-stage regression, the type I error is seen to be somewhat conservative, except when G is used as the instrument and there is genetic confounding (Figs 3 and 4, panels F and I; Fig S3, panel A). In contrast, when the first-stage uncertainty is not taken into account, the correct type I error (corresponding to the chosen *α* level) is generally observed (Figs 3 and 4, panels D, E, G, H; Figs S1 and S2, panels G and J; Fig S3, panel J), again except when G is used as the instrument and there is genetic confounding (Figs 3 and 4, panels F and I; Fig S3, panel G). This conservative behaviour of ‘standard’ instrumental variable regression-based MR seen in these simulations may relate to our use of a relatively weak instrument; we found that increasing the magnitude of the regression coefficient *β_GX_*used to generate the simulated data resulted in generally well-calibrated type I error.

When MR Steiger is used, the type I error and power are both reduced compared to MR (resulting in a conservative test, provided a valid instrument is available) due to the extra condition that a variable must pass a *p*-value threshold to be selected as a valid instrument, whereas in MR this is already assumed. Under model 3, MR Steiger with G used as a possible instrument can show inflated type 1 error (Fig 3, panel I), while for MR Steiger with Z used as a possible instrument, both the type I error and the power are generally zero due to Z never actually being chosen as a valid instrument for X in any of the 1000 simulation replicates (Figs 3 and 4, panels C, F, I; Fig S6, panels A, C, D). This behaviour of MR Steiger never actually choosing the proposed instrument is also seen under models 1 and 2, when Z is used as a possible instrument and we wish to test from X to Y (Figs 3 and 4, panels D, E, G, H; Figs S4 and S5, panels A and C), or when G is used as a possible instrument and we wish to test from Y to X (Figs S4 and S5, panels B and D).

For BN, we find the power is generally higher when both G and Z are used together rather than using either alone (Figs 3 and 4, panels A and B). Under model 1, the probability of making an incorrect inference is very low when testing Y to X when there is actually an effect from X to Y (Fig S4, panel B) or vice versa (Figs 3 and 4, panel G), and zero when there is no effect at all (Figs 3 and 4, panel D; Fig S4, panels C and D). Under model 2, there is a small chance of making an incorrect inference when testing X to Y (and no such effect exists,) particularly under strong confounding when Z is included as a possible explanatory variable (Fig 4, panels E and H). For model 3, there is a fairly large chance of making an incorrect inference when testing X to Y, if in fact there is an effect from Y to X (Fig 3, panel I), or vice versa (Fig S6, panel B), when there is weak confounding and G is included as a possible explanatory variable.

#### Receiver operating characteristic (ROC) curves

Figs 5 and 6 show receiver operating characteristic (ROC) curves for MR/MR Steiger and BN for testing the relationship X to Y under simulation models 1-3, with assumed effect size *β_XY_* or *β_Y X_* = 0.5. The results in the middle and right hand plots (models 2 and 3) in Fig 5 involve weak confounding, while those in Fig 6 involve strong confounding. The curves are constructed with respect to testing for a causal effect from X to Y, where the curves for true/false positives are constructed by gradually relaxing the detection threshold (based on type I error *α* for MR, or posterior probability of existence of an arrow for BN) used. For the top plots (panels A-C), false positives on the x-axis are counted using simulations when there is no effect (*β_XY_* = 0), while true positives on the y-axis are counted under simulations when there is an effect (*β_XY_* = 0.5). For the bottom plots (panels D-F), the false positive rate is calculated in a slightly different way, by simulating from a model where there is a causal effect from Y to X. Overall, the ROC curves are most appealing for BN, showing a generally higher power for a given type error rate. Similar conclusions apply when comparing BN with MR’ and St’ (performed using two-stage least squares regression without accounting for the uncertainty of the predicted values in the first-stage regression), see Figs S10 and S11.

**Fig 5.**
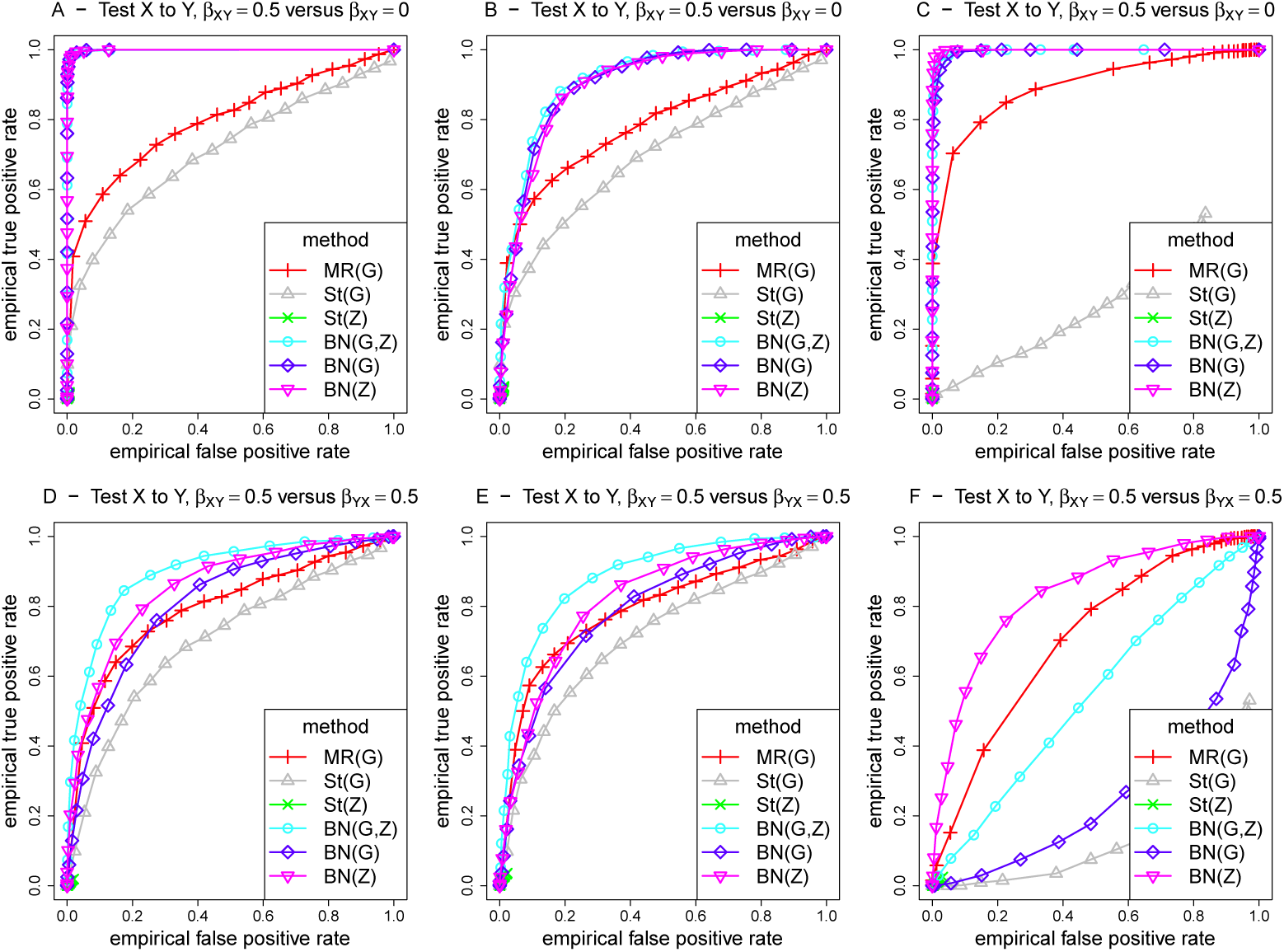
ROC curves for different methods for detecting an edge from X to Y, under different generating scenarios that include weak confounding. MR and St denote MR and MR Steiger respectively, performed using instrumental variable regression which takes into account the uncertainty of the predicted values in the first-stage regression to calculate the MR *p*-values. Left hand plots (A, D, G) are generated under model 1 (no confounding), middle plots (B, E, H) are generated under model 2 (non-genetic confounding), and right hand plots (C, F, I) are generated under model 3 (genetic confounding). For the top plots (panels A-C), false positives on the x-axis are counted using simulations when there is no effect (*β_XY_*= 0), while for the bottom plots (panels D-F), the false positive rate is calculated by simulating from a model where there is a causal effect from Y to X.

**Fig 6.**
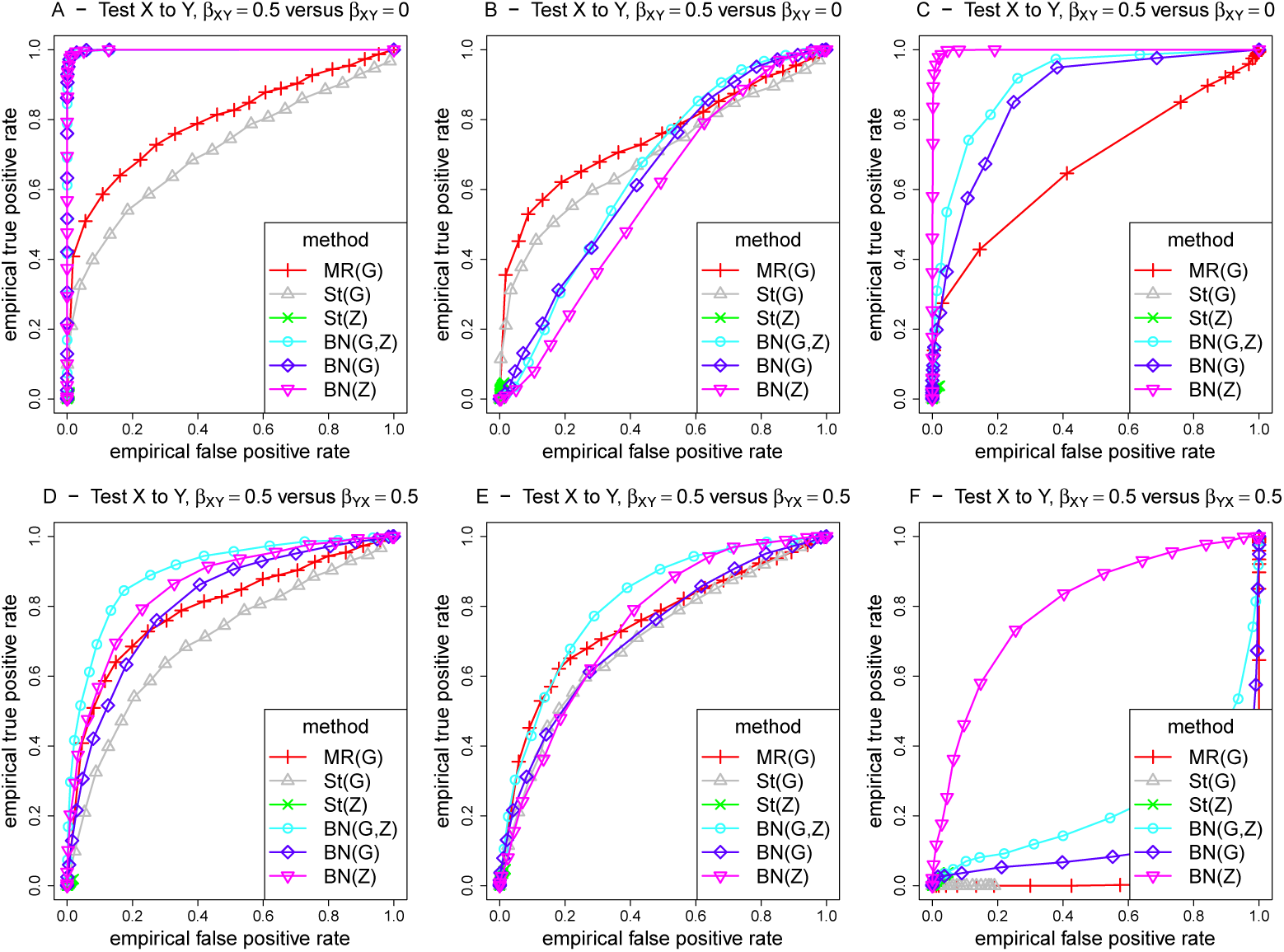
ROC curves for different methods for detecting an edge from X to Y, under different generating scenarios that include strong confounding. MR and St denote MR and MR Steiger respectively, performed using instrumental variable regression which takes into account the uncertainty of the predicted values in the first-stage regression to calculate the MR *p*-values. Left hand plots (A, D, G) are generated under model 1 (no confounding), middle plots (B, E, H) are generated under model 2 (non-genetic confounding), and right hand plots (C, F, I) are generated under model 3 (genetic confounding). For the top plots (panels A-C), false positives on the x-axis are counted using simulations when there is no effect (*β_XY_*= 0), while for the bottom plots (panels D-F), the false positive rate is calculated by simulating from a model where there is a causal effect from Y to X.

#### BN inference on direction of causality

To illustrate the ability of BN to infer the direction of causality, Fig S12 shows box plots of the probability estimates of X being causal on Y (top row) and of Y being causal on X (bottom row) given by BN, for data simulated under model 1 where X was causal on Y. As the true effect size *β_XY_* increases, the probability of correctly (top row) detecting an X to Y effect increases, while when *β_XY_* is zero, the probability of incorrectly detecting this effect is near zero. When both G and Z are included in the analysis (panel A) the probability estimates are higher than when only one of these is included (panels B and C), illustrating the advantage of using the extra information from both variables. In addition, as the true effect size, *β_XY_*, increases, the probability of falsely (bottom row) detecting a Y to X effect decreases, with the lowest probability seen when both G and Z are included in the analysis (panel D), again illustrating the advantage of using the extra information from both variables. Fig S13 shows the BN box plots for data simulated under model 2, the non-genetic-confounding model. Here the probabilities are all closer to 0.5, illustrating that BN does not work quite as well in this scenario as under model 1. In particular, when there are no effects between X and Y, the probabilities are much further from zero, with a mean well above 0. Fig S14 shows the BN box plots for data simulated under model 3, the genetic-confounding model. This shows better estimates of near zero when there is no effect from X to Y. However, for the analysis with only G included, it can be seen the probability estimates approach 0.5 as the effect size increases, rather than increasing to above 0.5 as occurred for models 1 and 2.

An overall summary of the performance of the methods based on Simulation Study 1 is given in Table 2. Naturally, these conclusions pertain only to the (relatively simple) set of scenarios that we have investigated here. In particular, we have not investigated the presence of both genetic and non-genetic confounding, nor have we considered situations such as when *β_CX_* and *β_CY_*have opposite signs. The robustness of BN to such types of confounding would be an interesting topic for future investigation. One can certainly envisage many other more complex simulation scenarios that would be interesting to explore, but are beyond the scope of this initial investigation. More complicated scenarios involving more complex networks of variables including extreme pleiotropy are, however, investigated in Simulation Study 3 (below).

**Table 2.**
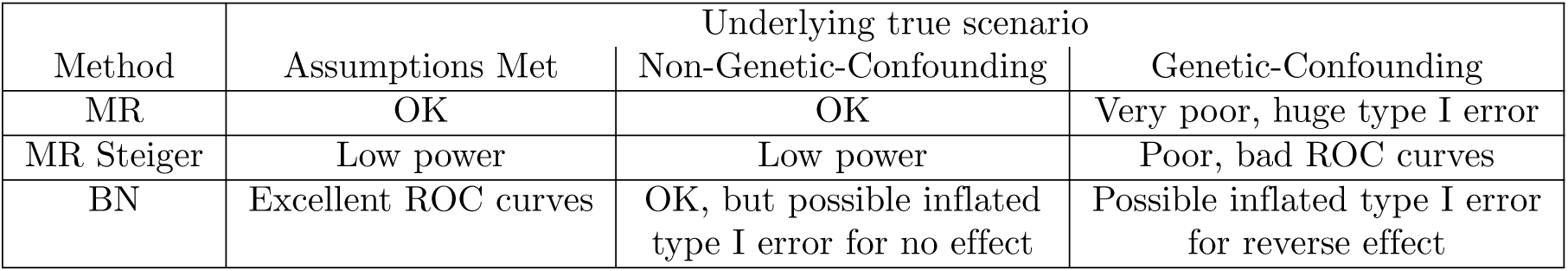
Performance of MR, MR Steiger and BN on the simulated quantitative trait data

### Simulation Study 2

We also investigated the utility of BNs for performing causal inference in a binary trait setting. Discrete binary data was simulated for 5000 individuals using four models considered in recent work by Shih et al. [55], in the context of quantifying the effects of alcohol consumption and high alanine transaminase levels on hepatocellular carcinoma. Data were simulated according to Fig 7, under four different scenarios with generating model parameter values as listed in Table 3. Fig 8 shows the average BN inferred when genetic variable Q was constrained to have no parents. For scenarios A and B, the only edges detected are those starting at (directed away from) Q, as the generating parameters for these causal relationships outweigh any others (the estimates of which do not meet the strength threshold to be plotted). The generating parameters under the other two models (panels C and D) are more evenly balanced, and this is reflected by most edges appearing in the average network with a probability strength of at least 0.4.

**Fig 7.**
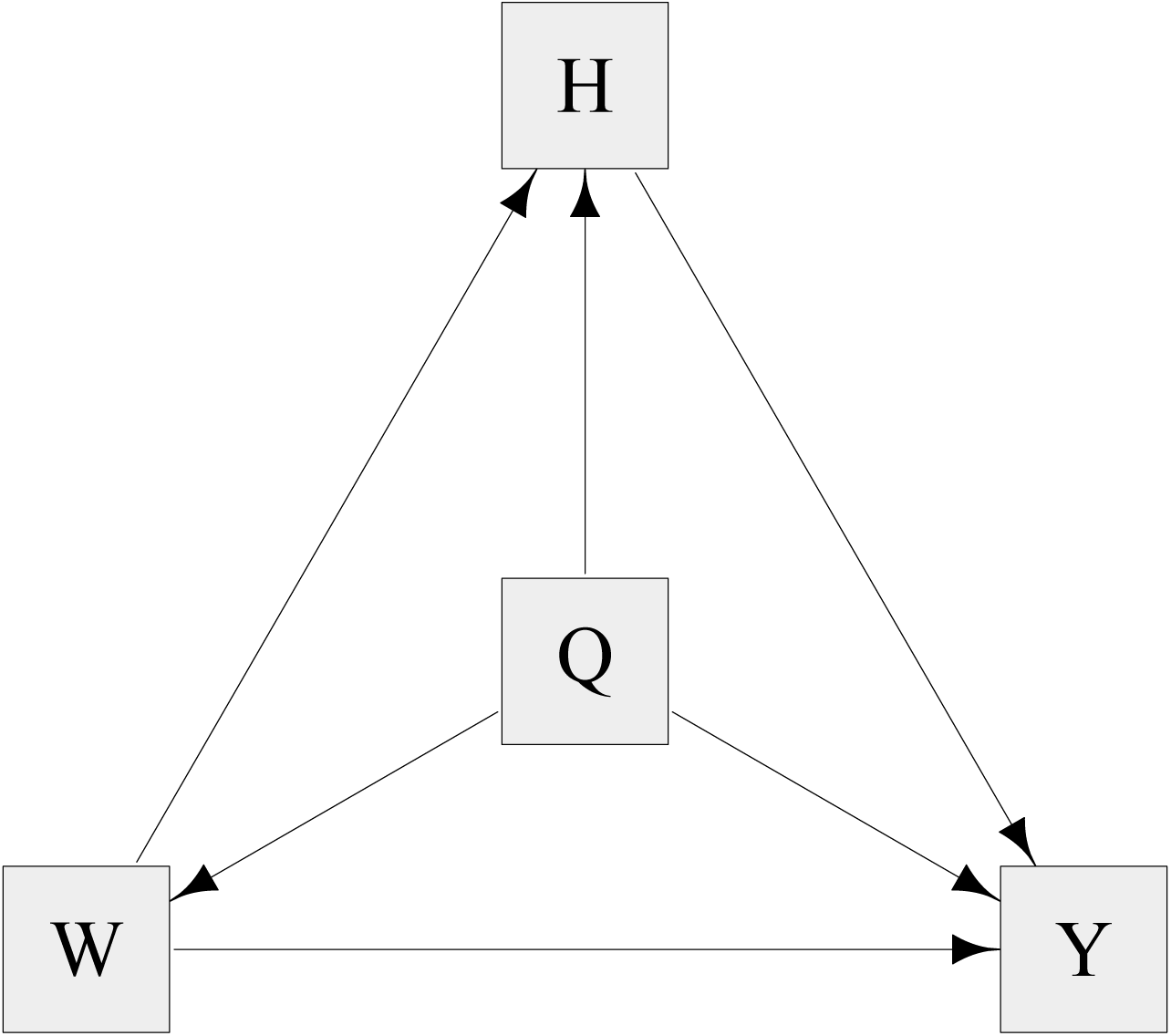
Graph of the simulation model used for Simulation Study 2 for four different parameter scenarios as described by Shih et al. [55]. The data simulated consisted of four binary variables: Q, representing a gene; W, representing high alcohol; H, representing high alanine transaminase; and the outcome variable, Y, representing hepatocellular carcinoma.

**Fig 8.**
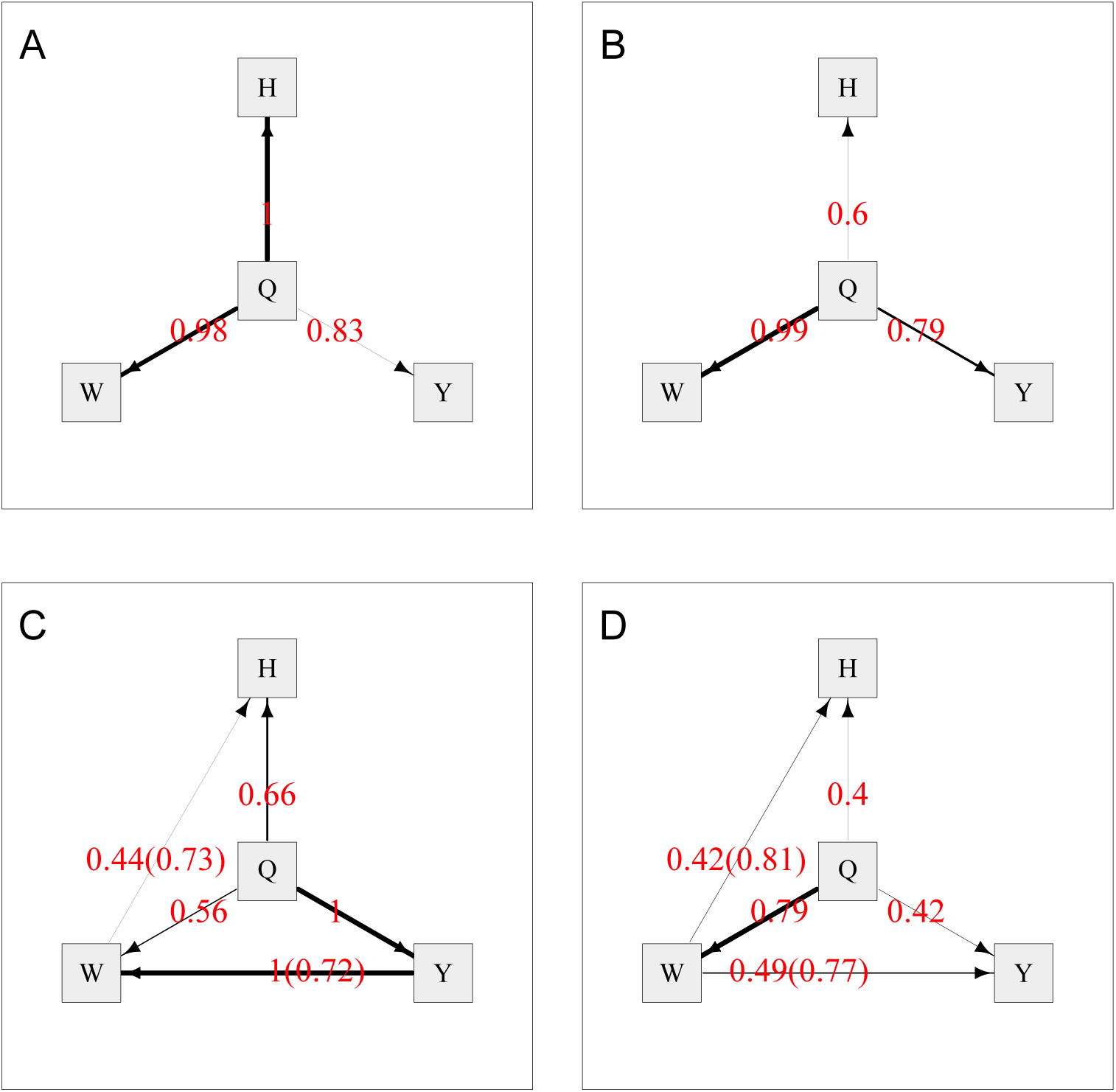
Average Bayesian networks for each of the four scenarios (A–D) used for the simulated binary data. The red numbers indicate the probability of existence of an edge, and the numbers in brackets indicate the probability of the edge operating in direction shown, given that it exists. The thickness of the edges indicates their strength (probability of existence). G is constrained to have no parents.

**Table 3.** The parameter values for each scenario used to simulate discrete binary data. Models are described in detail by Shih et. al. [55].

| Scenario | Frequency of Q=1 | Parameters influencing Y |  |  |  | Parameters influencing H |  |  | Parameters influencing W |  |
| --- | --- | --- | --- | --- | --- | --- | --- | --- | --- | --- |
| | | $\beta_0$ | $\beta_q$<br>(Q to Y) | $\beta_W$<br>(W to Y) | $\beta_H$<br>(H to Y) | $\alpha_0$ | $\alpha_q$<br>(Q to H) | $\alpha_W$<br>(W to H) | $\delta_0$ | $\delta_q$<br>(Q to W) |
| A | 0.49 | 0.2 | 0.3 | 0.25 | 0.15 | 0.2 | 0.4 | 0.2 | 0.1 | 0.3 |
| B | 0.49 | -0.9 | -0.3 | -0.1 | 0.2 | 0.0 | 0.2 | 0.2 | 0.2 | 0.3 |
| C | 0.49 | -1.0 | -1.0 | -0.8 | 0.2 | 0.0 | 0.2 | 0.2 | 0.2 | 0.3 |
| D | 0.49 | 0.2 | 0.2 | 0.2 | 0.1 | 0.1 | 0.2 | 0.2 | 0.1 | 0.2 |

Fig 8C shows an edge from Y to W with direction probability 0.72, which is in the opposite direction from how it was simulated. We explain this by noting that the two edges between Q and Y, and between Y and W, are much stronger than other edges in the model, each with probability strength 1, and so are always inferred to be in the model. The weak relationship between Q to W can be modelled by an additional edge from Q to W, but can also be modelled via the edges from Q to Y to W, which has the advantage of using 2 edges rather than 3 edges to model the entire system of relations between Q, Y and W. Therefore, although incorrect, this model was sometimes chosen as the best model on account of the fact that the BIC used in the **bnlearn** algorithm penalizes the number of edges in the network.

Fig 9 shows the average BN for scenarios A–D when the model is fitted with the extra constraint that Y (representing the outcome, hepatocellular carcinoma) has no child nodes. The results are similar to Fig 8 except that panel C now has the edge from W to Y in the correct direction (as implied by the extra constraint), and the edge from Q to W is much stronger since it is now the best way to model the causal relationship between Q and Y. This highlights the fact that constraints (if known) should be added to provide better causal inference as well as to improve computational efficiency.

**Fig 9.**
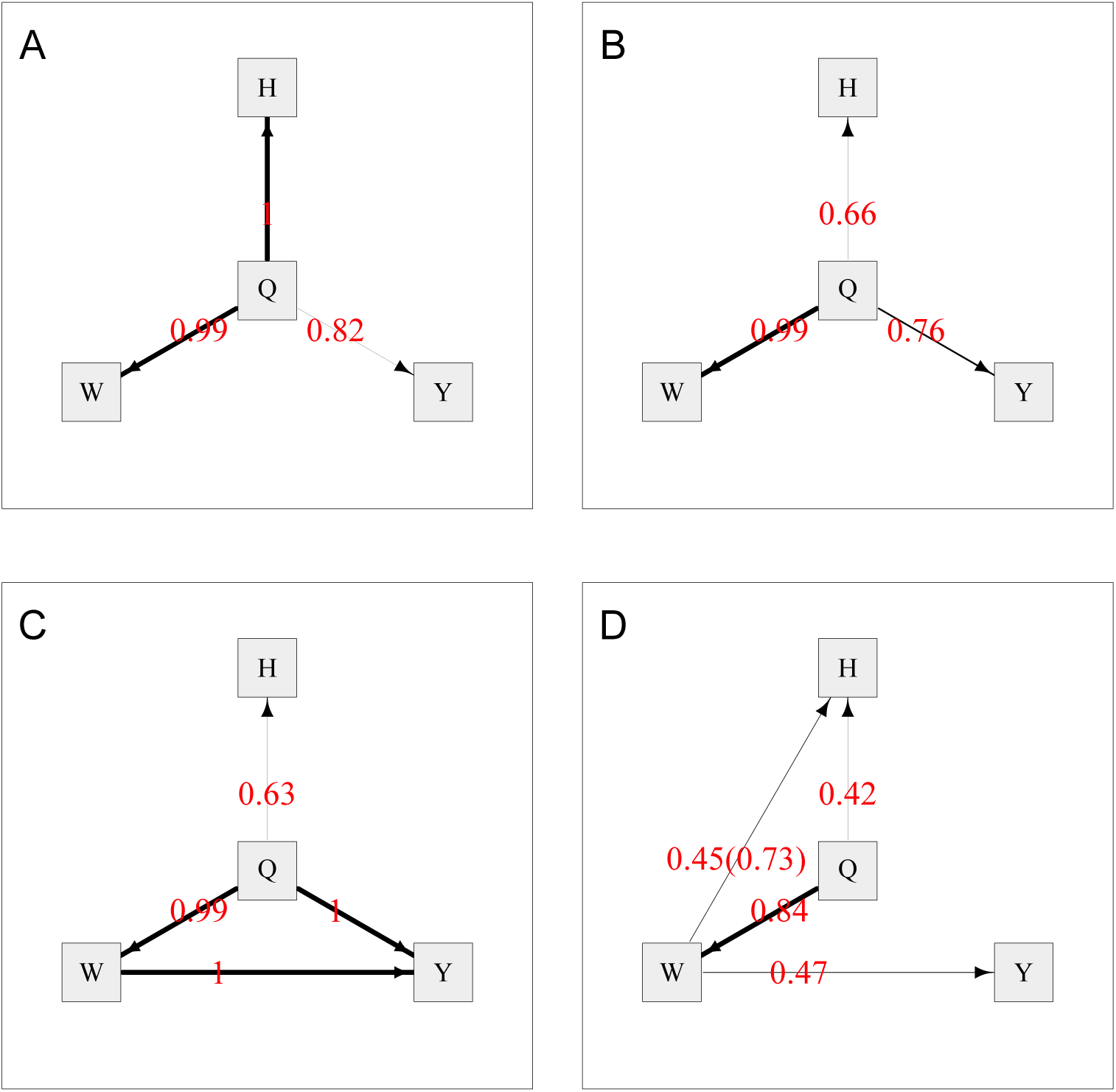
Average Bayesian networks for each of the four scenarios (A–D) used for the simulated binary data. The red numbers indicate the probability of existence of an edge, and the numbers in brackets indicate the probability of the edge operating in direction shown, given that it exists. The thickness of the edges indicates their strength (probability of existence). G is constrained to have no parents and Y is constrained to have no children.

### Simulation Study 3

We also carried out a simulation study involving more complex networks of variables including extreme pleiotropy (see Methods). This included 4 metabolites (Fig 10, left hand panels) simulated to have no effects, 4 metabolites (Fig 10, middle panels) simulated to have a causal effect on the outcome Y, and 4 metabolites (Fig 10, right hand panels) with a reverse effect (so that Y influenced the metabolite). We applied the recently proposed SMUT [48] method, along with BN, MR and a recent MR extension (MR-BMA) [25].

**Fig 10.**
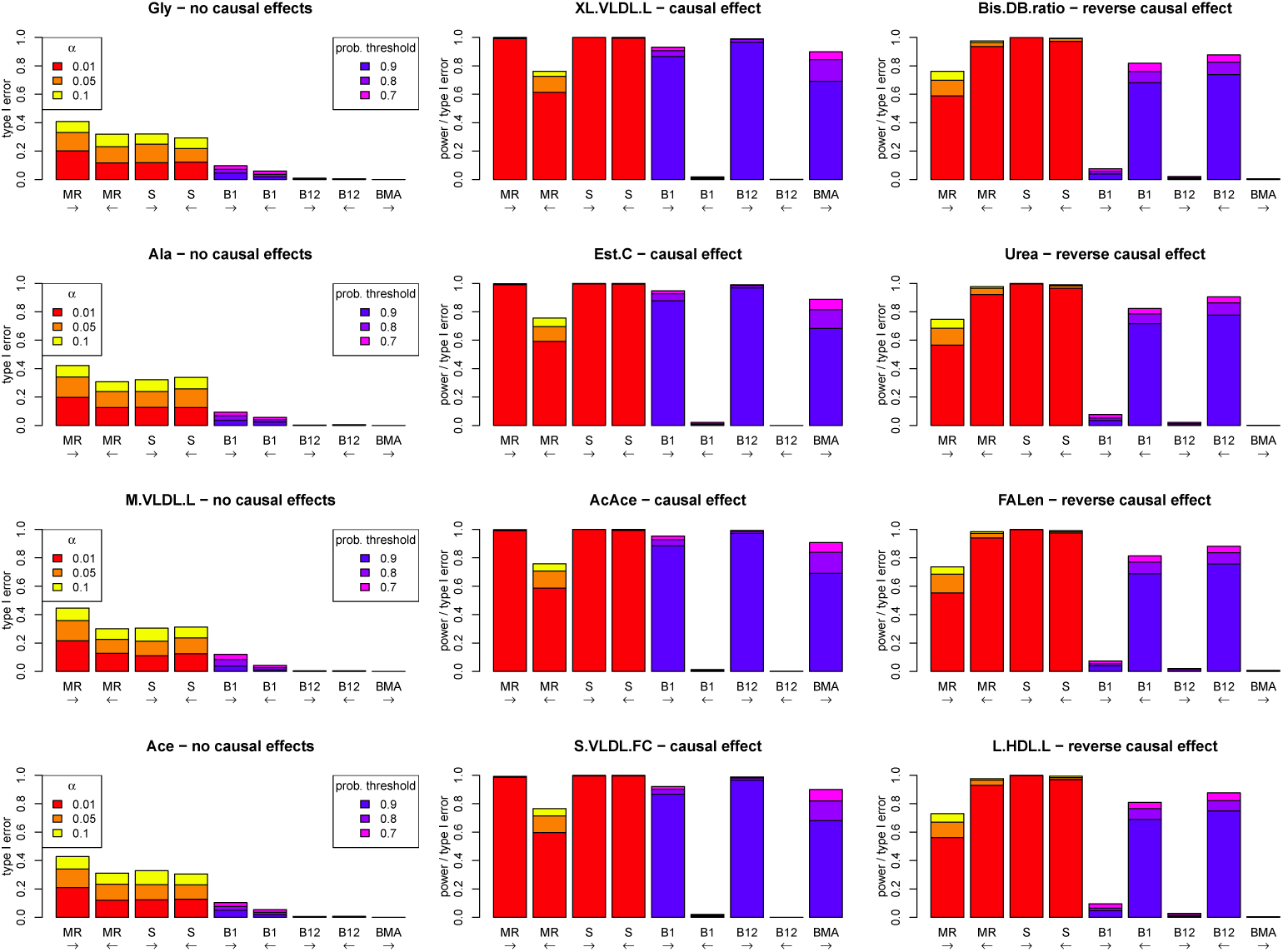
Performance (power and type I error) of different methods under a simulation model with 12 metabolites, an outcome Y, 150 SNPs affecting the metabolites, 75 other SNPs affecting Y, and 9775 SNPs with no effect. Four metabolites (middle panels) have a causal effect on Y, four metabolites (right hand panels) have a reverse causal effect from Y to the metabolite, and four metabolites (left hand panels) have no effects to Y in any direction. The left-to-right arrows show tests for a causal effect from the metabolite to Y, and right-to-left arrows show tests from Y to one of the metabolites. MR: Mendelian randomization using an allele score as an instrumental variable for one of the metabolites or Y. S: SMUT, using SNPs as random effect variables for one of the metabolites or Y. B1: Bayesian network consisting of one metabolite, Y and the two corresponding allele score variables. B12: Bayesian network consisting of all 12 metabolites, Y and all corresponding allele score variables. BMA: multivariable MR based on Bayesian model averaging (MR-BMA)

Fig 10 shows the powers and type I errors of MR, SMUT, BN using one metabolite risk factor (B1), BN using all 12 metabolite risk factors (B12) and MR-BMA, for testing the relationship between each metabolite and Y. MR, SMUT and B1 tested the relationship between each metabolite and Y separately (ignoring any information from other metabolites), whereas MR-BMA and B12 tested the relationships between Y and all 12 metabolites in one analysis. MR, B1 and B12 were used to test for a causal effect between metabolites and Y in either direction while MR-BMA could only be used to test from the metabolites to Y. The methods required different approaches to handle the 10,000 SNPs that were potentially causal on the metabolites or Y: for MR, B1 and B12 a weighted allele score was constructed (using SNPs passing a *p*-value threshold of *p <* 5 × 10^−6^), while SMUT and MR-BMA used a subset of SNPs passing a *p*-value threshold of *p <* 5 × 10^−6^ with any metabolite.

For MR and SMUT, we see high power to detect a true causal relationship when it is present (Fig 10, middle panels), however there is very high inflated type I error when the effect is in the opposite direction (Fig 10, right hand panels). There is also inflated type I error when there are no effects at all (Fig 10, left hand panels). This is due to the instrumental variable assumptions being violated, similar to what was seen previously for MR in Simulation Study 1.

For BN with only one risk factor (B1), there is very high power to detect a causal relationship in the correct direction when it is present, due to the use of genetic instruments for both the metabolite and Y. The type I error when there is a reverse effect is also very low, despite our previous observations (in Simulation Study 1) suggesting potential inflated type I error when there is genetic confounding and a reverse effect. This may be due to the effect sizes of the SNPs directly affecting the metabolite and those affecting Y being quite different, meaning that the complex nature of the confounding does not interfere too much with inference here (although clearly the same robustness is not seen for MR).

For BN with all 12 risk factors (B12), there is very high power to detect causal relationships when they are present, even higher than with B1, most probably due to the complementary manner in which all of the data is used together to better resolve the direction of edges. The type I errors are also much lower than B1 for the same reason. The powers are higher than for MR-BMA, probably due to the fact that the SNPs that affect Y are modelled in BN but are not accounted for in the MR-BMA analysis.

MR-BMA shows very high power when there is an effect and very low type I error when there is no effect or a reverse effect. Although MR-BMA is powerful (with low type I error) at successfully selecting only those metabolites that affect Y, its power is slightly lower than that of BN, and, unlike BN, it cannot detect a reverse effect from Y to the metabolites.

We also analysed these data using the recently-developed latent causal variable (LCV) method [29], which uses all 10,000 SNPs to test whether the “genetic causality proportion” (GCP) (the extent to which part or all of the genetic component of one trait is causal for another) differs from 0. However, in comparison to the other methods considered, the results (Fig S15) were rather poor, with a very low rejection rate for the LCV test of GCP= 0 when there is a causal relationship between the metabolite and Y in either direction. The LCV tests of GCP= −1 and GCP= 1 did, however, show good power to (correctly) reject the hypotheses that one variable was fully causal of another.

## Discussion

There is a growing interest in the use of causal inference methods in genetic epidemiology. With “omics” data becoming increasingly available in large cohort studies, we now have potential to uncover novel predictive and prognostic markers at the molecular level, variation in which may correlate with disease status even in its early stages. To make use of such markers, it is crucial to evaluate the extent to which any observed associations are due to causal relationships between the variables. For more than a decade, MR has been a popular approach used to strengthen or undermine a hypothesized causal relationship identified from epidemiological studies. However, conventional MR may not be readily applicable to infer simultaneous causal relationships in large-scale “omics” data because it generally deals with only one potential cause and one potential effect at a time. A naive application of MR, such as testing a causal effect of each “omics” variable on the disease outcome one at a time, may violate the no-pleiotropy assumption. Thus, recent developments in MR [19–26, 56] have focussed on considering several instruments and/or mediators simultaneously, in principle better accounting for horizontal pleiotropy. However, in most of these approaches, an underlying hypothesized graphical structure representing the relationships between variables must be assumed (rather than being learned from the data).

Here we propose Bayesian network analysis with the inclusion of genetic anchors to help orient relationships as a complementary approach for performing exploratory analysis of causal relationships in complex data. BN has many conceptual (and practical) similarities to MR, indeed BN in the context used here could be considered as an implementation of the basic logic of MR [4]. Both MR implemented through IV analysis and BN rely on assuming a specific graphical model (as well as making certain other assumptions) to carry out inference regarding the relationships between two or more variables of interest, and the inference obtained will be affected by any violation of the assumed relationships. In particular, unmeasured confounding is a problem for both methods, although MR does have the advantage of being robust to unmeasured confounding that only operates between the mediator and the outcome (i.e. non-genetic confounding), while the same is not necessarily the case for BN. However, there are also differences between the approaches, in particular the algorithmic details (and required calculations) for the two methods as performed in practice are quite different. BN naturally offers some additional flexibility in terms of exploring a (potentially large) number of different graphical models, a goal that could be considered to have motivated the development of MR extensions such as bi-directional MR and MR Steiger. BN also allows an appealing quantification of the strength of evidence provided by the data for different relationships (and indeed for entire graphical models) via calculation of network scores and bootstrap estimates of posterior probabilities for specific relationships. This contrasts with the inference provided from MR which is generally presented in frequentist terms and is based on *p*-values (or effect estimates and their standard errors). Despite recent debate about the utility and potential for misinterpretation of *p*-values in scientific research [54], we note that *p*-values can still be considered as useful summaries of the compatibility between a data set and an underlying hypothesized model [54, 57]. While this interpretation is only strictly correct if the entire assumed data generating model (not just the lack of existence of the targeted effect) holds, genetic associations identified using this paradigm have generally proved highly reproducible and, in the human genetics literature, *p*-values remain the most commonly-used summary measures indicating the extent of evidence for potential associations [58]. Indeed, it is this very fact that underpins their potential utility for use in MR. Thus, while we concur with the opinion [54] that the use of absolute significance thresholds should be avoided (and we do not propose that any particular threshold should be considered as “correct”), we still consider *p*-values to be useful summary measures that may be used, as in our illustrative example, to inform the comparison of competing hypotheses, or, as in our simulation studies, as a heurstic to examine the relative performance of different methods (in terms of true and false detections of relationships) as the thresholds are varied.

Our motivating example involved a situation where two correlated risk factors were associated with the same genetic variant, and showed how BN could help to resolve their simultaneous potential causal effects. Our example illustrated that BN could infer causal relationships even in the absence of a genetic anchor for the risk factor, as long as a genetic anchor for the outcome is available. In principle, BN can be applied to more complicated data with much larger numbers of variables [46], as long as the conditional dependencies of the variables are graphically representable as DAGs. Thus BN complements MR in terms of allowing a deeper exploration of possible underlying causal structures, and in providing a natural way of including multiple instruments and/or mediators in the analysis, similar to the goal of a number of recent MR extensions [19–26, 56].

We conducted a series of simulation studies and showed that BN with two directional anchors outperformed bi-directional MR based on ROC curves of the true positive rate (i.e. power) for a fixed false positive rate (i.e. type I error). The behaviour of both approaches will depend on the sample size and the absolute value of the causal effect as well as on the presence of confounding; both BN and MR perform better as the true causal effect (or the sample size) increases. BN performs better with more directional anchors available, since they remove uncertainty in the model search and help orient the true causal directions between other variables. However, even when only a single directional anchor is available, BN performs as well as or better than MR, at least under the scenarios and parameter values considered here. In our study, the performance of both BN and MR was affected by genetic confounding but barely affected by non-genetic confounding. (However, note that non-genetic confounding can, in some circumstances, create serious bias, and MR has begun addressing this by carrying out between-sibling analyses which are protected from the common sources of this bias [59]). In models involving pleiotropic relationships, BN outperformed both MR and the recently-proposed MR-BMA method, as well as outperforming SMUT.

We also considered a recently-developed latent causal variable (LCV) method [29] which infers, for pairs of measured traits, the extent to which part or all of the genetic component of one trait is causal for the other. This method analyses one mediator variable and one outcome variable at any given time, making use of genetic data across the whole genome, rather than following the usual MR approach of selecting specific genetic variants to be used as instruments. However, the approach is conceptually rather different from the other methods considered here, and unlike the other methods, does not formally test whether the mediator is causally related to the outcome (or vice versa). Exploratory analysis (Fig S15) using data from Simulation Study 3 suggested that the LCV test of GCP= 0 was not well-powered in this scenario, possibly because the concept of a genetic causality proportion (GCP) was not directly encapsulated by our simulation model.

We are not the first to propose the utility of genetic anchors to help orient relationships between variables in the context of network analysis [43–45, 47]. However, to our knowledge, this is the first study to directly compare the performance of MR and BN in both real and simulated data. Previously, Ainsworth et al. [60] applied MR, BN and structural equation modelling to simple simulated data scenarios and noted that BN and structural equation modelling could offer potentially attractive alternative (or at least complementary) approaches to MR. Given that structural equation modelling and BN overlap in many situations (for example, if the graphical model is a DAG and the local distribution follows a normal distribution), this current study corroborates that suggestion. We show here that BN with both directional anchors has greater power than bi-directional MR when applied to the same data, and further report on scenarios where BN could be more easily applied than MR. These include data with multiple risk factors and/or data with no genetic variant for one of the risk factors available.

Like any analytical approach, BN has limitations. Its assumptions (required for valid causal inference) are easily violated. For example, modelling all possible causes and confounding factors of all variables in the data is usually impossible (although this limitation is shared with most other methods for causal inference). BN cannot explain a cyclic or feedback relationship among variables, whereas bi-directional MR can test this to some extent. The performance of BN is affected by the sample size and true causal effect sizes, and the posterior probability threshold that corresponds to any particular type 1 error rate is therefore hard to define. In addition, although not explored in our current study, BN is also more likely than MR to be adversely affected by measurement error [28]. Arguably the most serious limitation of BN is the fact that analysis is performed on individual level data, and the method is not readily extended to summary data (although this represents an interesting topic for future investigation). In contrast, MR approaches such as two-sample MR can utilize previously generated results, including those based on summary statistics, to make robust causal inference. Indeed, this is the predominant mode of MR analysis at present (although there may well be a move back to single sample individual participant data analysis in the future, given the availability of large-scale studies such as UK Biobank [61]). These latter two limitations (the sensitivity to measurement error and the requirement for individual level data) mean that BN will not replace MR, but instead offers a complementary approach, offering some advantages and some disadvantages. Both BN and MR rely on some fairly strong assumptions (such as the known graph for MR and the lack of latent confounders for BN) that should ideally be assessed in real data analysis. But, as BN and MR depend on different sets of assumptions, and ones that would not necessarily lead to distortion of the results to the same degree (or even in the same direction), findings from the two approaches could contribute to the triangulation of evidence regarding a particular causal question [62]. This would preferably be a pre-planned exercise, and where possible would include estimates from other approaches utilizing methods aimed at strengthened causal inference, all of which could be biased, but with orthogonal sources of bias affecting each [63].

There are some limitations to our current study. First, in our simulations, we consider only fairly simple scenarios with relatively small numbers of variables where both BN and MR can easily be applied (although we note that BN can readily be extended to utilize larger numbers of variables [43–46]). It is possible that BN may perform differently or worse in larger, more complex data sets. Thus, further studies on more complex real and simulated data (for example involving known biological or metabolic pathways) are required. In spite of these limitations, our study highlights the utility of BN as an appealing approach for performing causal inference in complex biological data sets that thus warrants further investigation.

## Materials and methods

### Mendelian Randomization

Mendelian randomization was performed using two-stage least squares linear regression [64]. The first linear regression used one or more genetic variables (either a single SNP, two SNPs or an allele score) as the explanatory variable(s) and the hypothesized risk factor as the response variable. The second linear regression used the predicted values of the risk factor for each individual from the first regression as the explanatory variable, and the outcome as response variable. We used instrumental variable regression using the function ivreg() from the R statistical software package AER [65], which takes into account the uncertainty of the predicted values in the first-stage regression to calculate the MR *p*-values. These *p*-values were used as a measure of evidence of a causal relationship. We also performed two-stage least squares regression using the lm() function in R, but without accounting for the uncertainty of the predicted values in the first-stage regression (denoted by MR’). We note that this method provides equivalent inference (identical *p*-values) to that obtained by simply regressing the outcome variable on the genetic variable(s).

We note that MR was originally [4] introduced as a general approach that uses the directionality from genetic variable to phenotype as the basic principle, but not with any particular analytical strategy (such as that suggested by its use here) in mind. In practice, the large majority of MR studies have attempted effect estimation and have used either two-stage least squares linear regression for analysis carried out within a single study sample, or two-sample MR based on summary statistics when utilizing data from two separate studies [15, 17] (which provides equivalent inference). This motivates our choice of two-stage least squares linear regression as reflecting the most commonly used analysis strategy, while also having the advantage of allowing direct comparison with BN (which, at least in its current implementations, requires access to individual level – rather than summary statistic level – data).

### Bayesian Networks

A variety of algorithms have been proposed for learning the structure and parameters of a Bayesian network. We considered two different algorithms, one motivated by the default algorithm implemented in the R package **deal** [53] and the other motivated by an option available in the R package **bnlearn** [39], which were implemented in C++ in our own software package, BayesNetty [66]. The **bnlearn** algorithm that we used corresponded to the Hill-Climbing score-based algorithm with Bayesian information criterion (BIC) score described on pages 52 and 106 of [39]. The BIC used to construct the network score has the structure of a penalized negative log-likelihood, meaning that models with lower BIC scores are considered to be a better fit, with the score for any hypothesized network inversely related to its posterior probability (calculated under specific distributional assumptions, namely that discrete nodes follow a multinomial distribution and continuous nodes a normal distribution, with distributional parameters determined by the values of the incoming parent nodes). The **deal** algorithm that we used involved specifying a prior network structure which is used, along with training data and default distributional assumptions, to generate an overall network score. For full details see [53]. For exploring the graph space, we used the same Hill-Climbing algorithm as was used with **bnlearn**. The **bnlearn** approach (as we demonstrate) was found to be more powerful and robust than **deal**. The **bnlearn** method is therefore the primary method used to generate the BN results presented in this article.

Networks were drawn using the igraph [67] R package. Average networks were calculated by bootstrapping the data with replacement 1000 times, and selecting the best-fit network for each replicate. The probability of an edge existing, and the probability of the edge being in a particular direction (given that it exists) were estimated by counting the proportion of times that such events occurred amongst the 1000 resulting best-fit bootstrap networks. For plotting the resulting average network, only edges that were considered sufficiently strong in the context of the current average network [39] were plotted.

### MR Steiger

In addition to MR and BN, we also considered a recently-proposed extension of MR known as MR Steiger [28]. This approach involves applying two tests which must both pass a *p*-value threshold in order to conclude a causal relationship between variables X and Y: firstly a test to decide if a genetic variable G is most suitable as an IV for variable X or Y, then a standard MR test using G as an instrument to test either the relationship X to Y, or the relationship Y to X.

### Multivariable MR based on Bayesian Model Averaging

We also considered a recently-developed extension to multivariable MR [19, 68] termed “multivariable MR based on Bayesian model averaging” (MR-BMA) [25]. MR-BMA, like original multivariable MR, is basically an extension of standard MR to model not one, but multiple, risk factors on an outcome, thus accounting for measured pleiotropy. MR-BMA aims to address the problem of selecting, from many potential causal risk factors, those that are most useful for one outcome variable, using Mendelian randomization principles. The method is based on inverse-variance weighted (IVW) linear regression in a two-sample framework, where the associations between genetic factors and the outcome (tested in sample 1) are regressed on the genetic associations with all the risk factors (tested in sample 2) in a multivariable regression approach.

### Latent Causal Variable method

We also considered the recently-developed latent causal variable (LCV) method [29] which infers, for pairs of measured traits, the extent to which part or all of the genetic component of one trait is causal for the other. This method makes use of genetic data across the whole genome, rather than following the usual MR approach of selecting specific genetic variants to be used as instruments. The method tests a newly-defined quantity between two traits, the “genetic causality proportion” (GCP), where large (positive or negative) values of GCP imply that interventions on one trait are likely to affect the other, suggesting (without specifically testing) that one trait may itself be causal on the other. Formally, the GCP test performs a two-sided test of the null hypothesis that the GCP= 0. The software also produces *p*-values for “full causality” between the two traits in either direction, testing the null hypothesis that GPC= 1 (implying trait 1 causes trait 2) or that GPC= −1 (implying trait 2 causes trait 1), respectively. The underlying graphical model used to motivate the LCV method actually corresponds to a model in which an (unmeasured) latent variable is the causal variable for both measured traits. One could therefore argue that demonstration of such an effect suggests that it is actually the latent variable that should be intervened upon, rather than one of the traits, if one wishes to bring about a corresponding change in the value of the other trait.

### Multi-SNP Mediation Intersection-Union Test

We also applied a recently-proposed multi-SNP mediation intersection-union test known as SMUT [48]. SMUT tests the joint mediation effects of multiple (potentially correlated) genetic variants on a trait through a single mediator, effectively generating a hypothesis test for mediation but with a multivariate genetic exposure. SMUT adopts the classical mediation framework, takes a frequentist approach, and relies on individual level data, treating the mediator effect as fixed and the effects of multiple SNPs upon the mediator as random.

### Illustrative Example: fatty acids and BMI

The data analysed here consisted of 5654 twins with measurements of BMI, the two metabolites EPA and DGLA, and 42 SNPs. For the purposes of the current analysis we used all twins and treated them as independent (i.e. ignoring pairwise clustering due to twin relationships; this will overestimate the statistical significance (nominal value) of any observed associations, but we anticipate that this phenomenon should affect all the methods evaluated equally). Three SNPs were used directly as IVs: rs174556 for EPA, and rs968567 and rs6498540 for DGLA. The other 39 SNPs known [51, 52] to be associated with BMI were combined into a weighted allele score variable (i.e. the number of effect alleles for each individual were counted for each the 39 SNPs, and these were summed, weighting by their previously estimated [51, 52] regression coefficient). The EPA and DGLA data were pre-processed using the same approach used by Shin et al. [50]. Specifically, the metabolite values were log transformed, the resulting transformed variables were adjusted for study day using linear regression, and the residuals from the regression were then transformed to the quantiles of a standard normal distribution to produce the final metabolite variables used for analysis. The genetic variables (rs174556, rs968567, rs6498540 and the weighted allele score) were used as instruments in MR to explore causal relationships between the metabolites and BMI. Relationships between all variables were also explored via BN, with the directional constraint that arrows were constrained to come out from (rather than go into) any nodes corresponding to genetic variables.

### Simulation Study 1

We also investigated the performance of BN and MR via computer simulations of a quantitative trait. Data were simulated for two continuous variables (X and Y), together with a genetic instrument G (coded as 0, 1, 2, mimicking a SNP) and a continuous instrumental variable Z (mimicking a SNP allele count). Data were simulated for 2500 individuals under three different generating models (shown in Fig 1), using a variety of values for the regression coefficients (the *β*s). These models cover a variety of plausible scenarios in terms of potential confounders (*C* and *S*). In each case, the direction of causality goes from X to Y.

For all three simulation models, the following analyses were implemented:

1. MR(G): Test the relationship X to Y using MR with G used as an IV for X.
2. MR(Z): Test the relationship Y to X using MR with Z used as an IV for Y.
3. St(G): Test the relationship X to Y (or Y to X) using MR Steiger with G used as an IV for X (or Y).
4. St(Z): Test the relationship X to Y (or Y to X) using MR Steiger with Z used as an IV for X (or Y).
5. BN(G,Z): Perform BN with variables X, Y, G and Z.
6. BN(G): Perform BN with variables X, Y, G only.
7. BN(Z): Perform BN with variables X, Y, Z only.

We also performed MR and MR Steiger without accounting for the uncertainty of the predicted values from the first-stage regression (denoted as MR’ and St’ respectively). As noted above, this provides equivalent inference (identical p-values) to that obtained by simply regressing the outcome variable on the chosen IV.

For BN, G and Z were constrained to operate as instruments i.e. the direction of the arrows was constrained to come out from (rather than go into) these nodes. For the purpose of network fitting, all variables were treated as continuous, regardless of whether they actually followed a continuous or discrete distribution.

For MR and MR Steiger, powers and type I errors (based on 1000 simulation replicates) were calculated for different values of *β_XY_* (ranging from 0 to 0.5), with *α* thresholds of 0.01, 0.05 and 0.1 used to define detection of a relationship. For BN, powers and type I errors (based on 1000 simulation replicates) for testing X to Y and Y to X were calculated with *β_XY_* equal to either 0 or 0.5, with probability thresholds 0.7, 0.8 and 0.9 used to define detection of a relationship. As a further visualization of the performance of BN for different values of *β_XY_* ranging from 0 to 0.5, the estimated probabilities (based on the average bootstrap network) of an edge existing from X to Y and from Y to X were calculated for each of the 1000 simulation replicates, and the distributions plotted as box plots.

Receiver operating characteristic (ROC) curves (based on 1000 simulation replicates) for the detection of an edge existing from X to Y were generated by imposing either different *α* thresholds (for MR and MR Steiger) or different probability thresholds (for BN). As the relevant threshold is relaxed, the chance of a true positive detection of a relationship increases, but so does the chance of a false detection of a relationship. Evaluation of power (a true positive effect from X to Y) was performed by simulating data under models with *β_XY_* equal to 0.1, 0.3 and 0.5. Evaluation of type 1 error (a false positive effect from X to Y) was performed by simulating data (a) under a model with no effect (i.e. with *β_XY_* equal to 0), and (b) under a model with an effect in the ‘wrong’ direction (i.e. from Y to X, with *β_Y X_* equal to 0.1, 0.3 or 0.5).

### Simulation Study 2

We also investigated the utility of BNs for performing causal inference in a binary trait setting. Discrete binary data was simulated for 5000 individuals using four models considered in recent work by Shih et al. [55], in the context of quantifying the effects of alcohol consumption and high alanine transaminase levels on hepatocellular carcinoma. We used the same graph (Fig 7) and parameter settings (Table 3) used by Shih et al. [55]. The data simulated consisted of four binary variables: Q, a gene; W, high alcohol; H, high alanine transaminase; and the outcome variable, Y, representing hepatocellular carcinoma. The data were analysed using BN with the constraint that Q has no parent nodes, and then again with the extra constraint that Y has no child nodes. For the purpose of network fitting, all variables were treated as multimomial (binomial), reflecting the fact that they followed a discrete distribution.

### Simulation Study 3

We also carried out a simulation study involving more complex networks of variables, as considered by Zuber et al. [25] in their development of the “multivariable MR based on Bayesian model averaging” (MR-BMA) method. We simulated data in a very similar manner to Zuber et al. [25] and then applied MR-BMA, along with BN, MR, and SMUT. Data was simulated for 1000 individuals, using 1000 replicates (allowing us to determine powers and type I errors using *p*-value thresholds of 0.1, 0.05 and 0.01, or posterior probability thresholds of 0.7, 0.8 and 0.9, respectively).

To inform our simulation model, we used the same publicly available summarized data on genetic associations with risk factors derived from a recent metabolite GWAS [69] as used by Zuber et al. [25]. To avoid selection bias we took the same subset of 150 independent SNPs as Zuber et al. [25], that had been found to be associated with any of the three main lipid measurements (LDL-cholesterol, triglycerides or HDL-cholesterol) at a genome-wide level of significance (*p*-value *<* 5 × 10^−8^) in an external data set, namely a large meta-analysis by the Global Lipids Genetics Consortium [70].

Beta-coefficients and standard errors of genetic associations between the 150 SNPs and the 118 metabolites with available data were extracted from the metabolite GWAS [69], in order to allow us to retain the empirically observed relationships between SNPs and metabolites. The set of metabolites was reduced by excluding at random one from each pair of metabolites that had a genetic correlation (calculated using the beta-coefficients of the 150 SNPs) stronger than |*r*| *>* 0.99. From the resulting 92 metabolites, 12 were chosen at random to be used in our simulation study. Four of the metabolites were chosen to be used in the simulation model as null variables (with no effects on the outcome variable, Y), four were chosen to be used in the simulation model with a direct effect on Y, and the other four were chosen to be used in the simulation model with a reverse effect (from Y to the metabolites).

The data for the 150 SNPs were simulated using the allele frequencies given in by the Global Lipids Genetics Consortium [70], assuming Hardy-Weinberg equilibrium (HWE). The four metabolites with direct effects on Y were simulated conditional on the simulated data for the 150 SNPs (based on their corresponding beta-coefficients for association with metabolites). Y was then simulated based on these four metabolites and 75 randomly-chosen SNPs (with beta-coefficients derived from their relationship with a randomly discarded metabolite, Ile). The metabolites not directly affecting Y were simulated based on the 150 SNPs and (for the 4 metabolites where a reverse effect was present) on Y. Any causal effects between the metabolites and Y, or vice versa, were simulated using a beta-coefficient of value 0.3, and the standard error was set to 1. A further 9775 SNPs with no effects on any other variables were simulated assuming HWE using a minor allele frequency simulated from a uniform distribution between 0.01 and 0.5. This gave a final simulated data set consisting of 10,000 SNPs, 12 metabolites and one outcome variable, Y.

We performed MR between every individual metabolite and Y, as well as MR in the reverse direction to test if Y has a causal effect on each metabolite. Weighted allele score variables were used as instrumental variables and were re-constructed within each simulation replicate using SNPs passing a *p*-value threshold of *p <* 5 × 10^−6^ of association with the appropriate metabolite or with Y, using the estimated regression coefficients as weights. The same SNPs were also used as the genetic variants in SMUT (using the R package SMUT [48]), which was also performed between every individual metabolite and Y (or vice versa for reverse causal effects).

The MR-BMA test was performed using R code written by the MR-BMA authors and was designed to detect which risk factors for an outcome are causal. The test outputs marginal probabilities for each metabolite being causal on the outcome variable; these are the probabilities presented in the results. The SNPs were chosen for the MR-BMA test using a *p*-value threshold of *p <* 5 × 10^−6^ for association with any of the metabolites.

Average Bayesian networks were used to estimate the probabilities of causal effects between the metabolites and Y using the same instrumental variables as used by the MR tests. Bayesian network analyses were initially performed using only 4 variables for every metabolite: the metabolite itself, the outcome Y, and the 2 corresponding allele scores (one for the metabolite and one for Y). Subsequently we considered Bayesian network analyses that used all 12 metabolites, Y, and the 13 relevant allele score variables (for the 12 metabolites and Y) simultaneously. In all analyses, the allele score variables were constrained to operate as individual genetic instruments i.e. to have one and only one edge going from itself to the corresponding instrumented variable (either a metabolite or Y).

We also applied the LCV method proposed by O’Connor and Price [29]. The test was evaluated using the R code written by the LCV authors and uses only one metabolite and the outcome variable at any one time, together with all 10,000 SNPs.

### Quantification of data underlying plots and graphs

We quantified the data underlying the various plots and graphs resulting from our analyses in S1-S20 Spreadsheets.

## Acknowledgments

We thank Chin Yang Shapland for pointing out some errors in, and Vanessa Didelez for providing useful comments on, previous versions of this paper. We thank TwinsUK for granting us access to their data. TwinsUK is funded by the Wellcome Trust, Medical Research Council, European Union, the National Institute for Health Research (NIHR)-funded BioResource, Clinical Research Facility and Biomedical Research Centre based at Guy’s and St Thomas’ NHS Foundation Trust in partnership with King’s College London.

## Supporting information

**S1 Fig.**
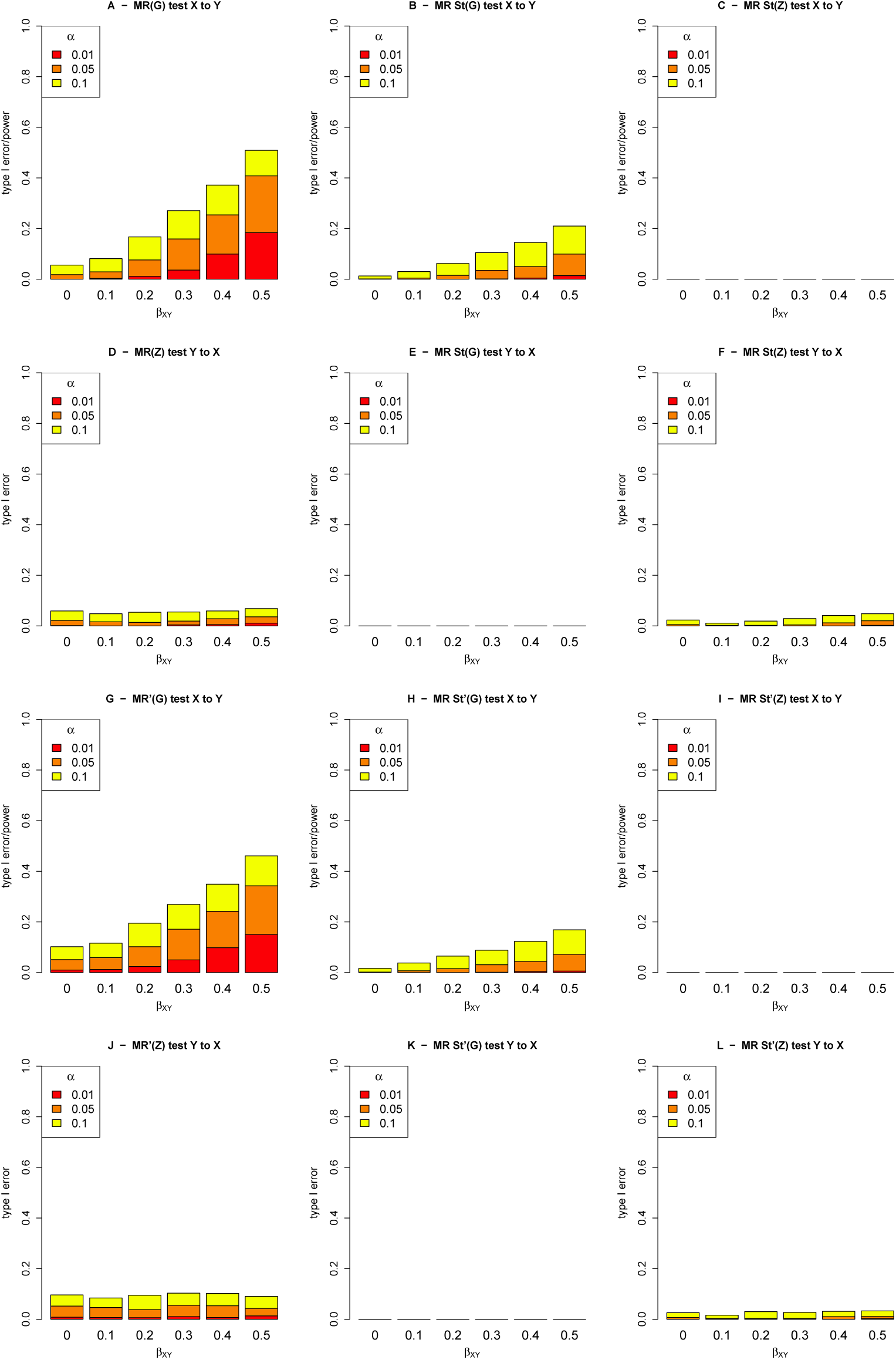
Power/type I error plots for MR and MR Steiger for data simulated under model 1. MR and St denote MR and MR Steiger respectively, performed using instrumental variable regression which takes into account the uncertainty of the predicted values in the first-stage regression to calculate the MR *p*-values. MR’ and St’ denote MR and MR Steiger respectively, performed using two-stage least squares regression without accounting for the uncertainty of the predicted values in the first-stage regression.

**S2 Fig.**
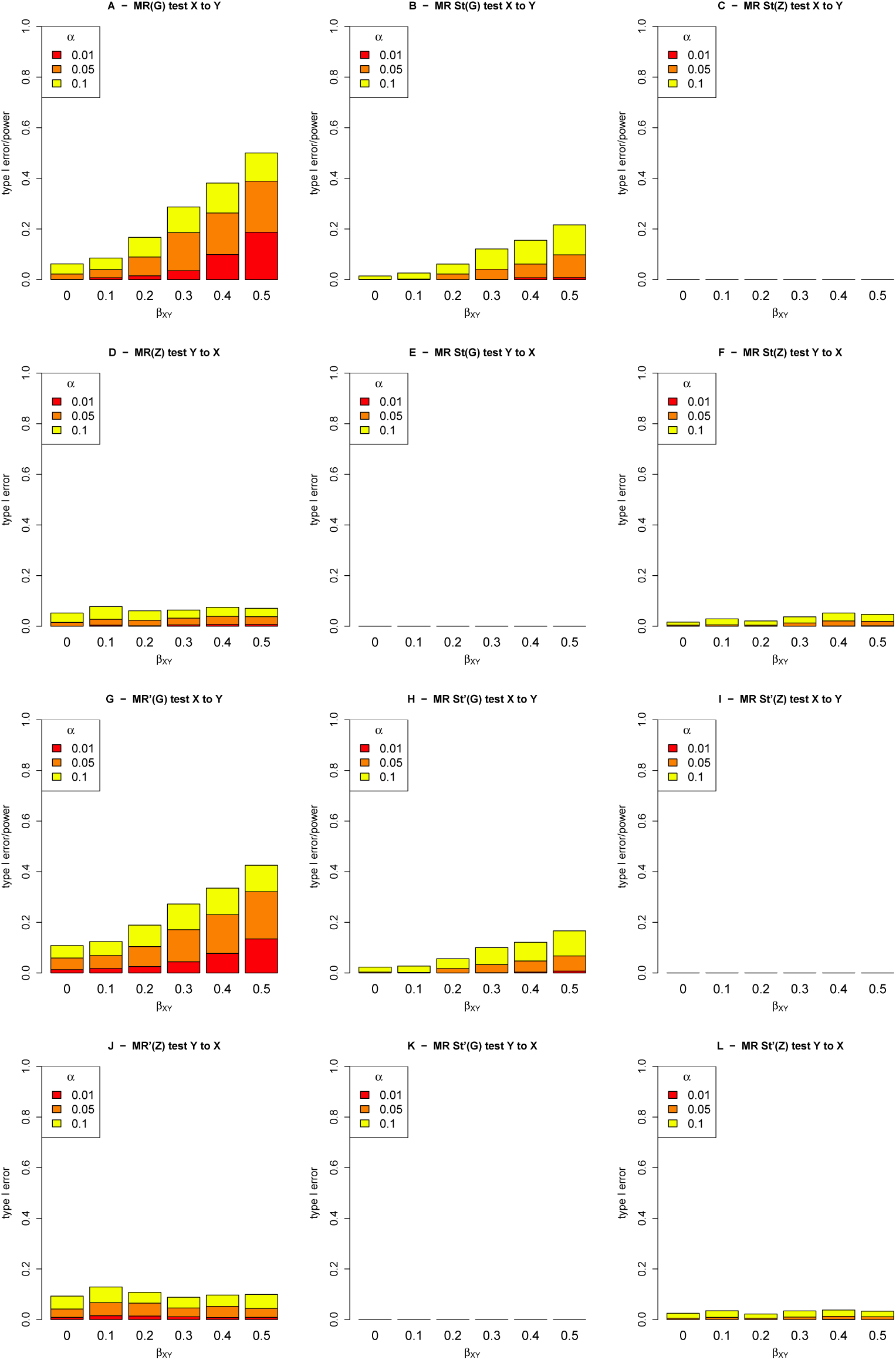
Power/type I error plots for MR and MR Steiger for data simulated under model 2. MR and St denote MR and MR Steiger respectively, performed using instrumental variable regression which takes into account the uncertainty of the predicted values in the first-stage regression to calculate the MR *p*-values. MR’ and St’ denote MR and MR Steiger respectively, performed using two-stage least squares regression without accounting for the uncertainty of the predicted values in the first-stage regression.

**S3 Fig.**
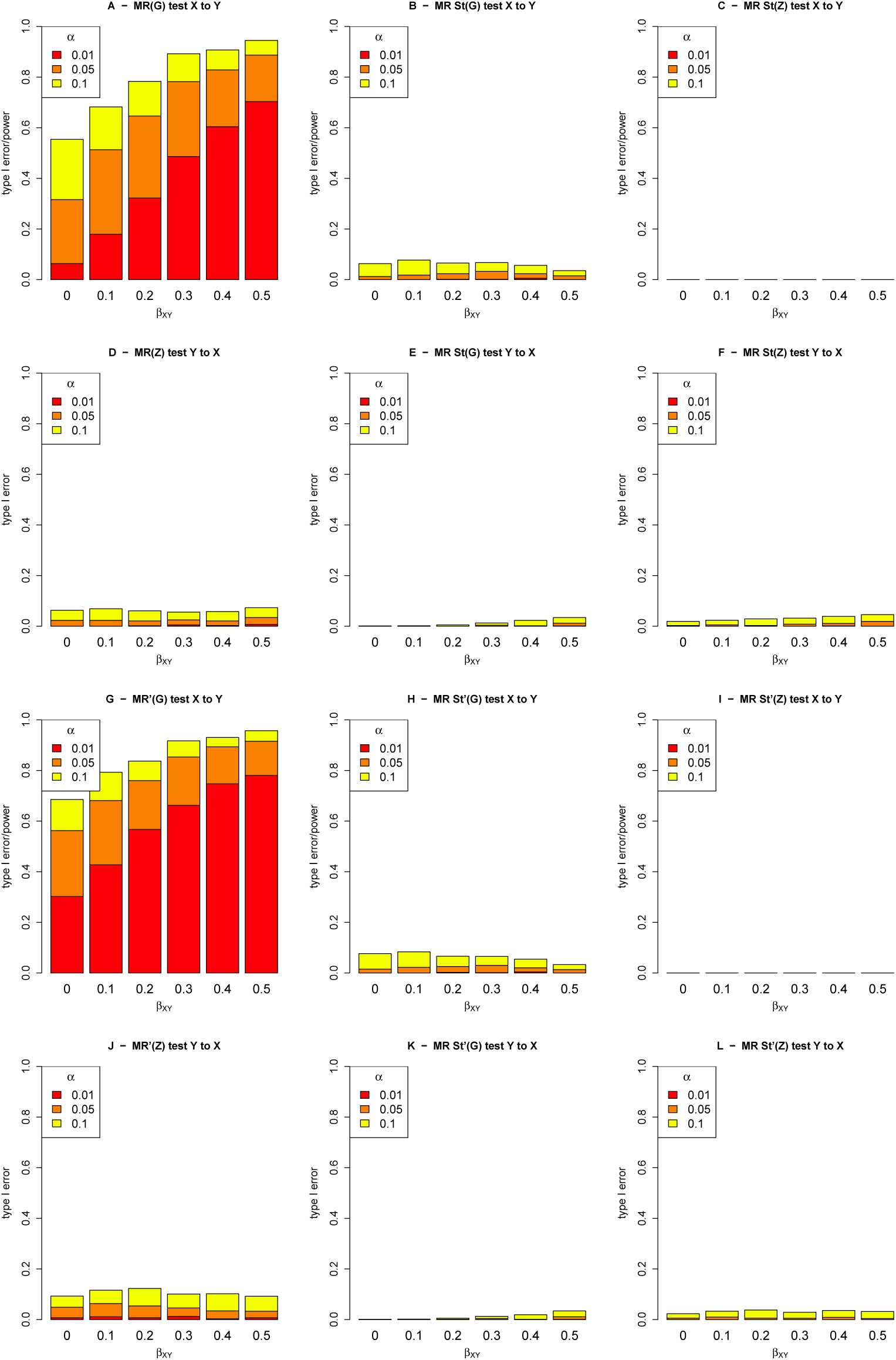
Power/type I error plots for MR and MR Steiger for data simulated under model 3. MR and St denote MR and MR Steiger respectively, performed using instrumental variable regression which takes into account the uncertainty of the predicted values in the first-stage regression to calculate the MR *p*-values. MR’ and St’ denote MR and MR Steiger respectively, performed using two-stage least squares regression without accounting for the uncertainty of the predicted values in the first-stage regression.

**S4 Fig.**
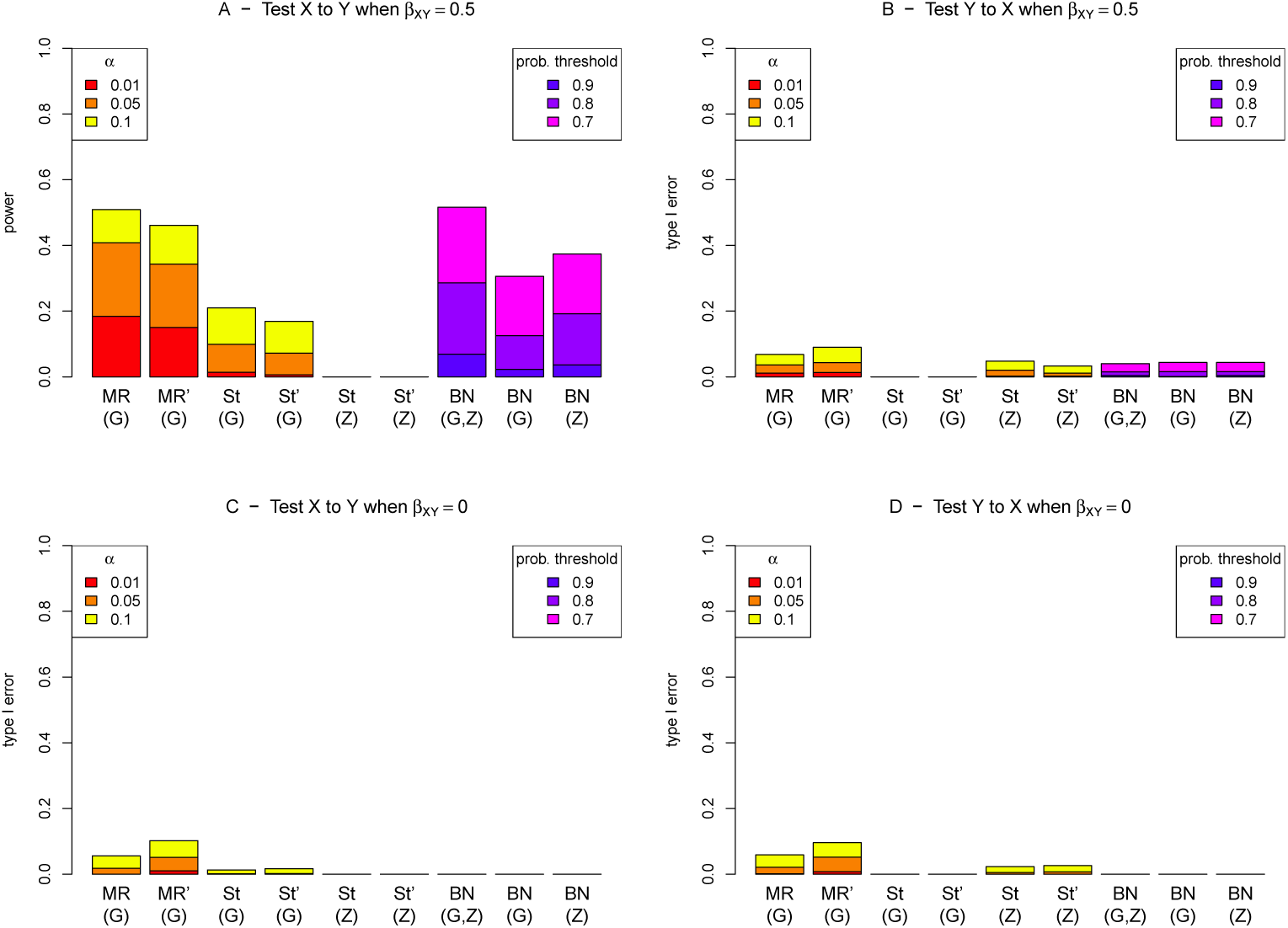
Power/type I error plots for MR, MR Steiger and BN (**bnlearn** algorithm) for data simulated under model 1. MR and St denote MR and MR Steiger respectively, performed using instrumental variable regression which takes into account the uncertainty of the predicted values in the first-stage regression to calculate the MR *p*-values. MR’ and St’ denote MR and MR Steiger respectively, performed using two-stage least squares regression without accounting for the uncertainty of the predicted values in the first-stage regression.

**S5 Fig.**
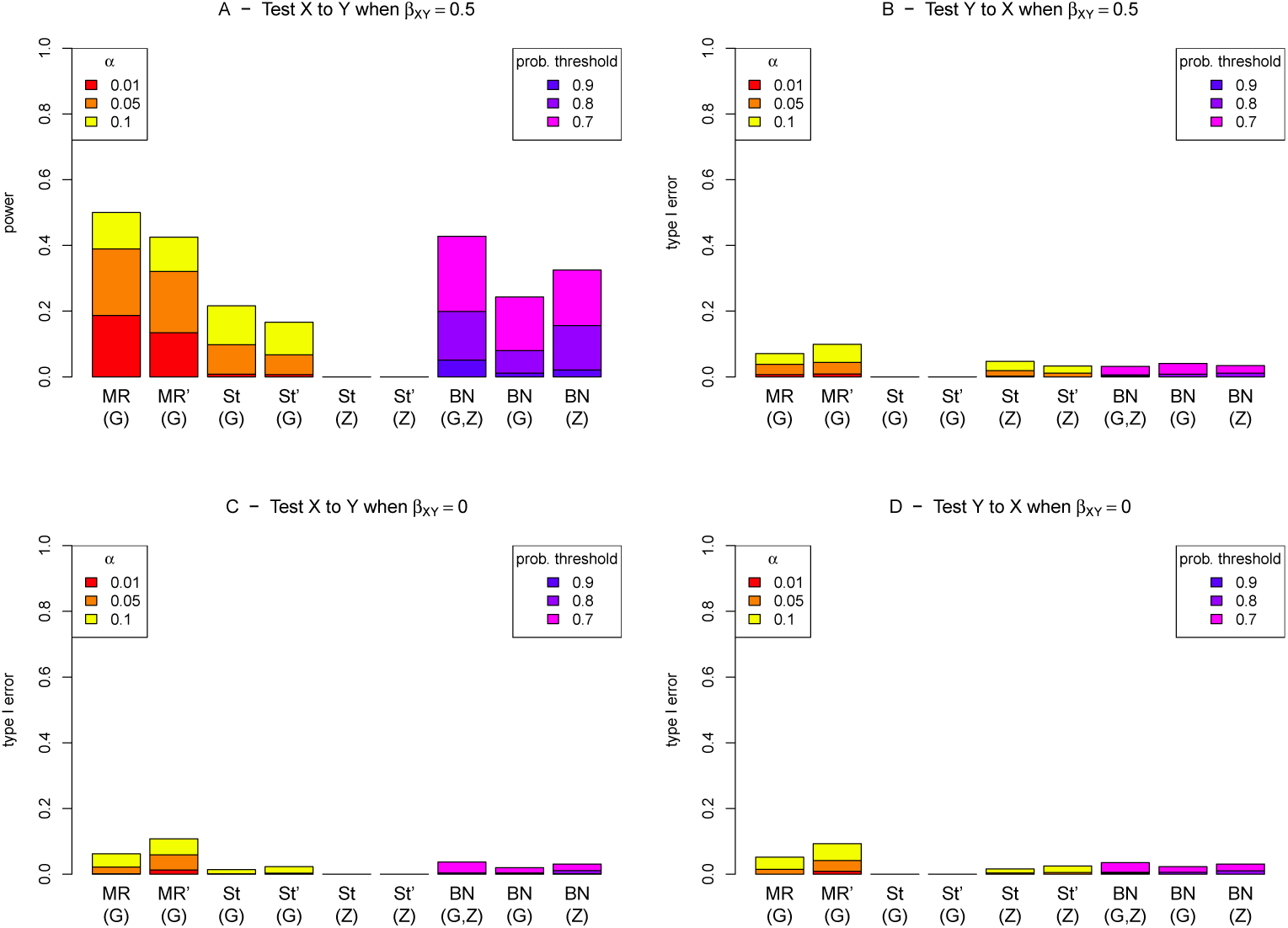
Power/type I error plots for MR, MR Steiger and BN (**bnlearn** algorithm) for data simulated under model 2. MR and St denote MR and MR Steiger respectively, performed using instrumental variable regression which takes into account the uncertainty of the predicted values in the first-stage regression to calculate the MR *p*-values. MR’ and St’ denote MR and MR Steiger respectively, performed using two-stage least squares regression without accounting for the uncertainty of the predicted values in the first-stage regression.

**S6 Fig.**
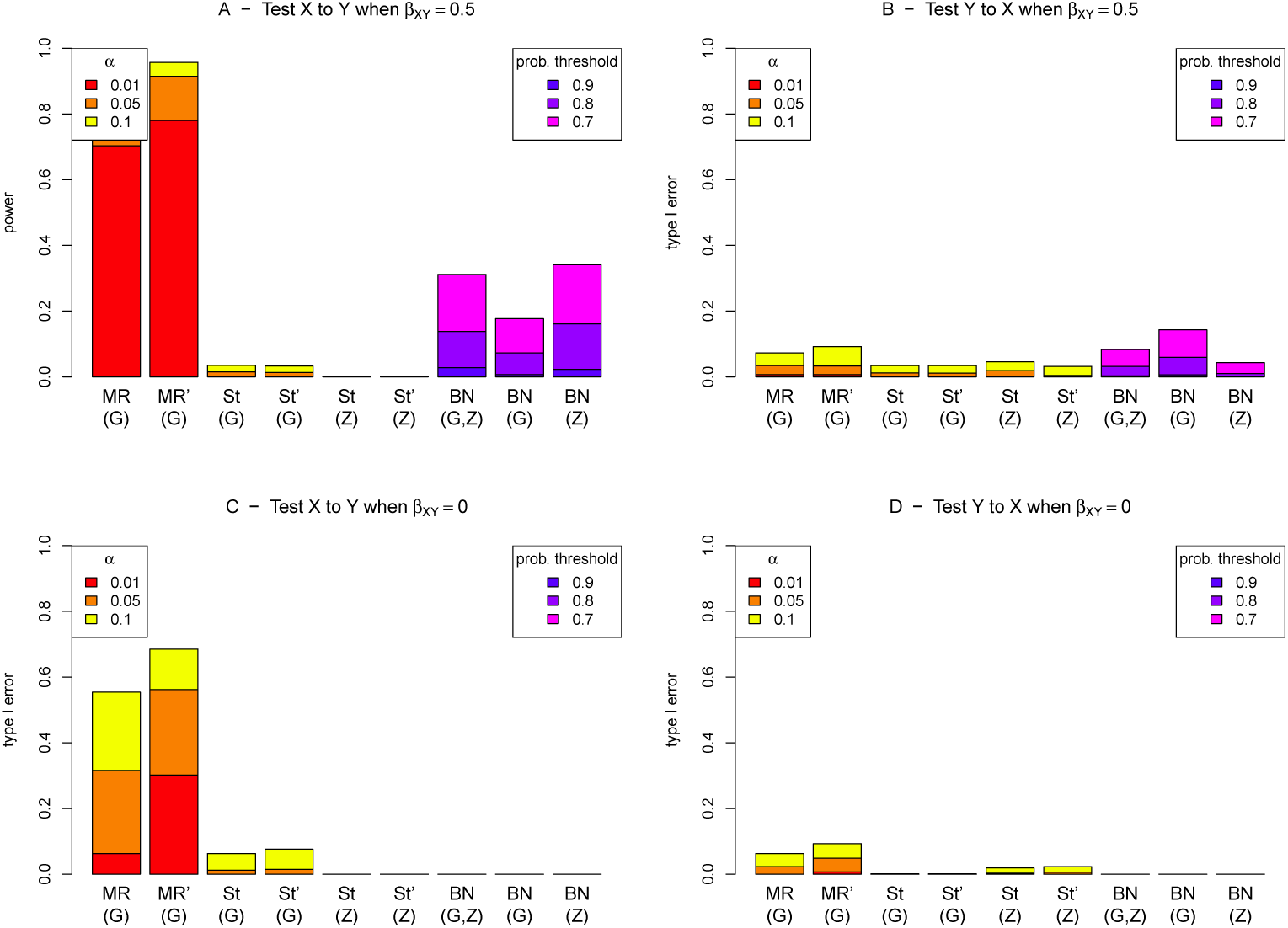
Power/type I error plots for MR, MR Steiger and BN (**bnlearn** algorithm) for data simulated under model 3. MR and St denote MR and MR Steiger respectively, performed using instrumental variable regression which takes into account the uncertainty of the predicted values in the first-stage regression to calculate the MR *p*-values. MR’ and St’ denote MR and MR Steiger respectively, performed using two-stage least squares regression without accounting for the uncertainty of the predicted values in the first-stage regression.

**S7 Fig.**
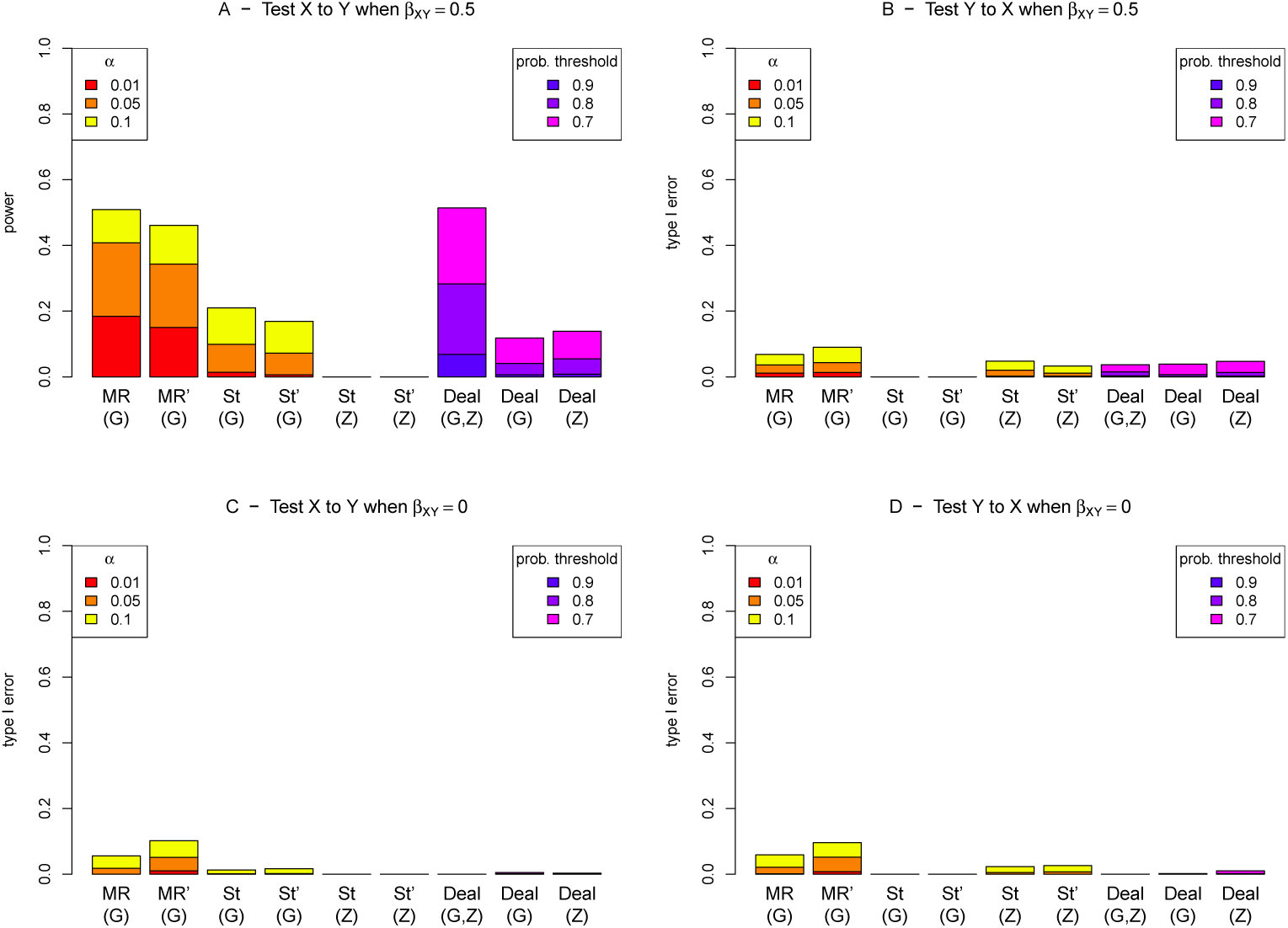
Power/type I error plots for MR, MR Steiger and BN (deal algorithm) for data simulated under model 1. MR and St denote MR and MR Steiger respectively, performed using instrumental variable regression which takes into account the uncertainty of the predicted values in the first-stage regression to calculate the MR *p*-values. MR’ and St’ denote MR and MR Steiger respectively, performed using two-stage least squares regression without accounting for the uncertainty of the predicted values in the first-stage regression.

**S8 Fig.**
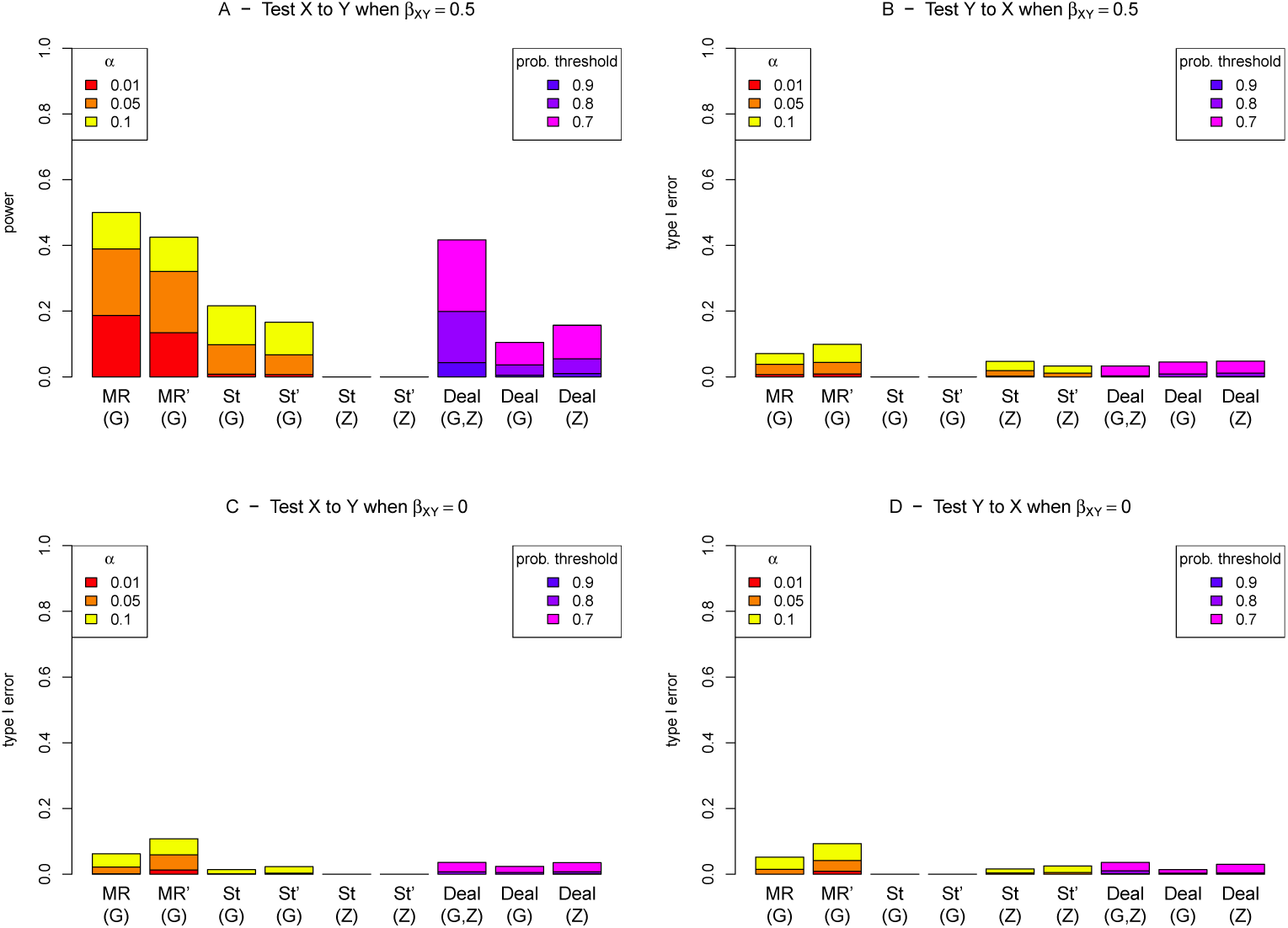
Power/type I error plots for MR, MR Steiger and BN (deal algorithm) for data simulated under model 2. MR and St denote MR and MR Steiger respectively, performed using instrumental variable regression which takes into account the uncertainty of the predicted values in the first-stage regression to calculate the MR *p*-values. MR’ and St’ denote MR and MR Steiger respectively, performed using two-stage least squares regression without accounting for the uncertainty of the predicted values in the first-stage regression.

**S9 Fig.**
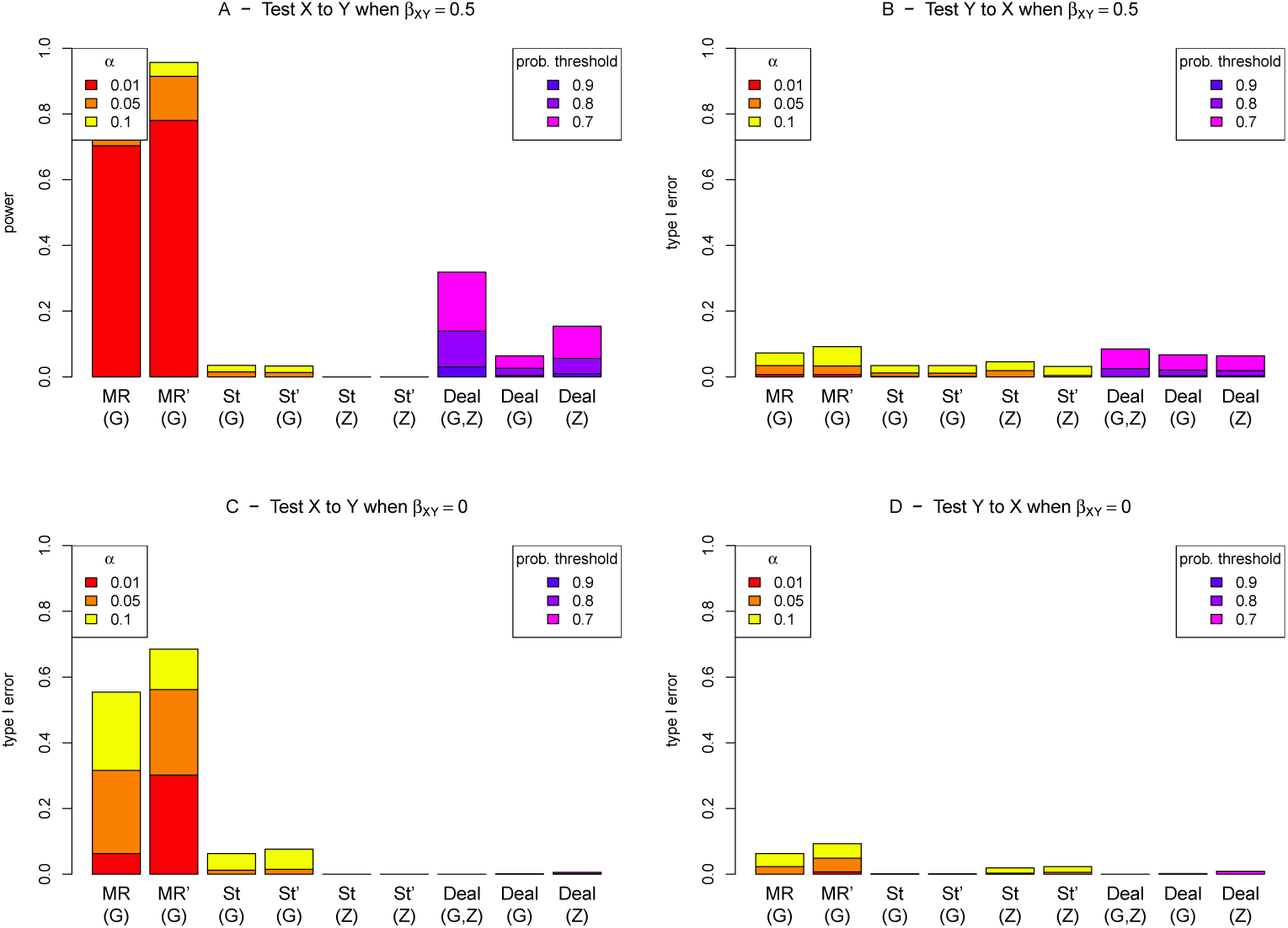
Power/type I error plots for MR, MR Steiger and BN (deal algorithm) for data simulated under model 3. MR and St denote MR and MR Steiger respectively, performed using instrumental variable regression which takes into account the uncertainty of the predicted values in the first-stage regression to calculate the MR *p*-values. MR’ and St’ denote MR and MR Steiger respectively, performed using two-stage least squares regression without accounting for the uncertainty of the predicted values in the first-stage regression.

**S10 Fig.**
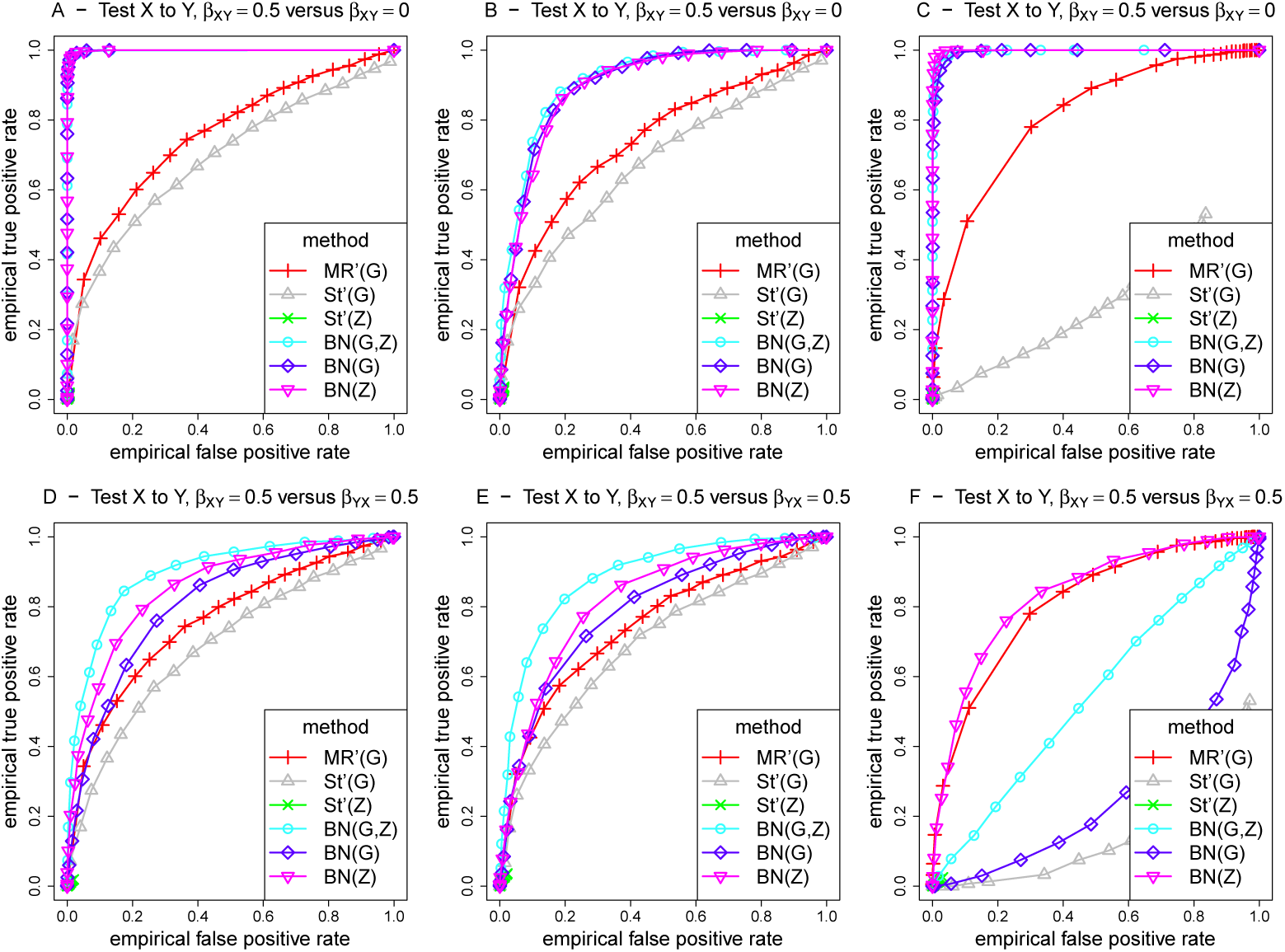
ROC curves for different methods for detecting an edge from X to Y, under different generating scenarios that include weak confounding. MR’ and St’ denote MR and MR Steiger respectively, performed two-stage least squares regression without accounting for the uncertainty of the predicted values in the first-stage regression. Left hand plots (A, D, G) are generated under model 1 (no confounding), middle plots (B, E, H) are generated under model 2 (non-genetic confounding), and right hand plots (C, F, I) are generated under model 3 (genetic confounding). For the top plots (panels A-C), false positives on the x-axis are counted using simulations when there is no effect (*β_XY_* = 0), while for the bottom plots (panels D-F), the false positive rate is calculated by simulating from a model where there is a causal effect from Y to X.

**S11 Fig.**
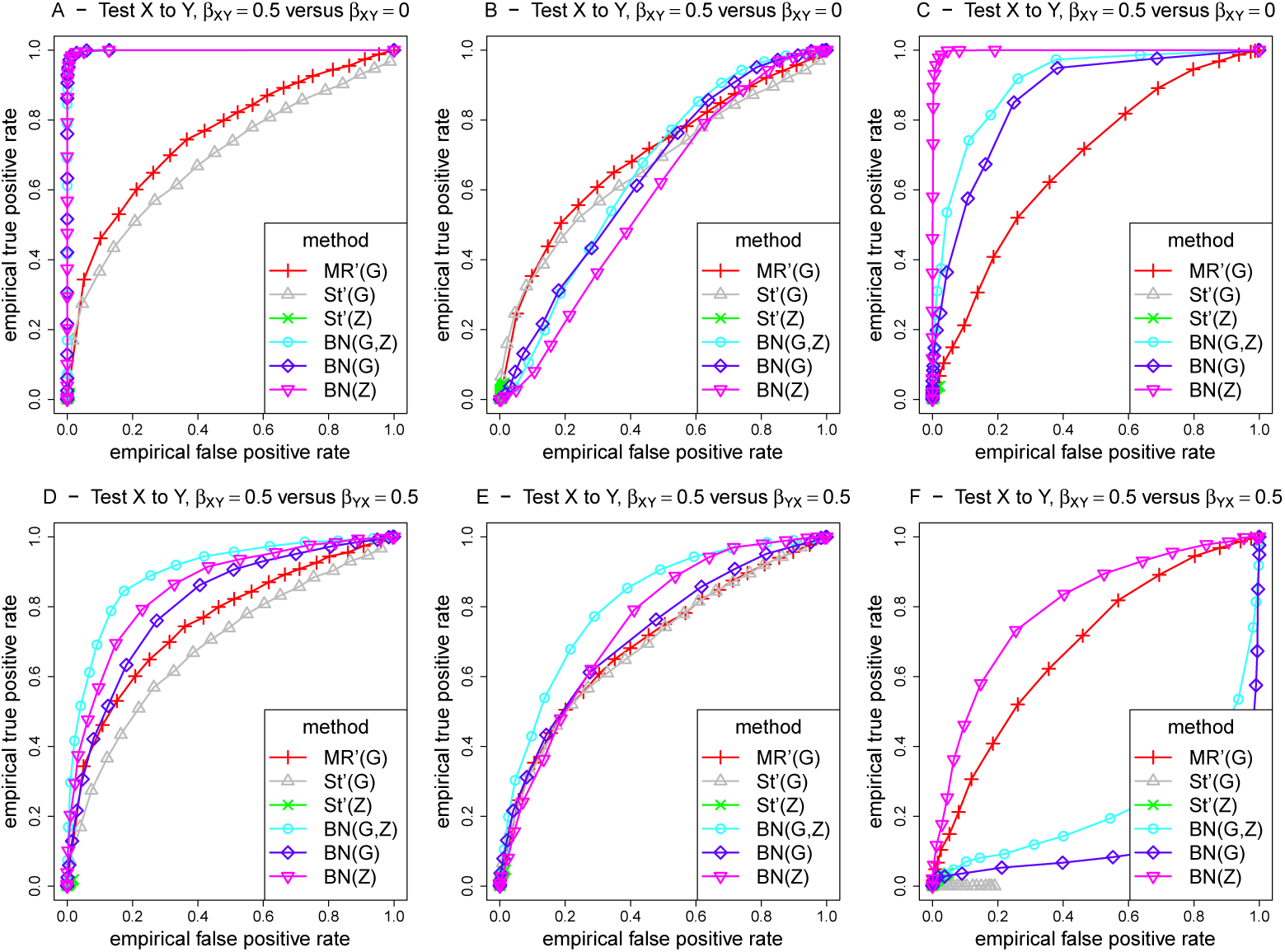
ROC curves for different methods for detecting an edge from X to Y, under different generating scenarios that include strong confounding. MR’ and St’ denote MR and MR Steiger respectively, performed two-stage least squares regression without accounting for the uncertainty of the predicted values in the first-stage regression. Left hand plots (A, D, G) are generated under model 1 (no confounding), middle plots (B, E, H) are generated under model 2 (non-genetic confounding), and right hand plots (C, F, I) are generated under model 3 (genetic confounding). For the top plots (panels A-C), false positives on the x-axis are counted using simulations when there is no effect (*β_XY_*= 0), while for the bottom plots (panels D-F), the false positive rate is calculated by simulating from a model where there is a causal effect from Y to X.

**S12 Fig.**
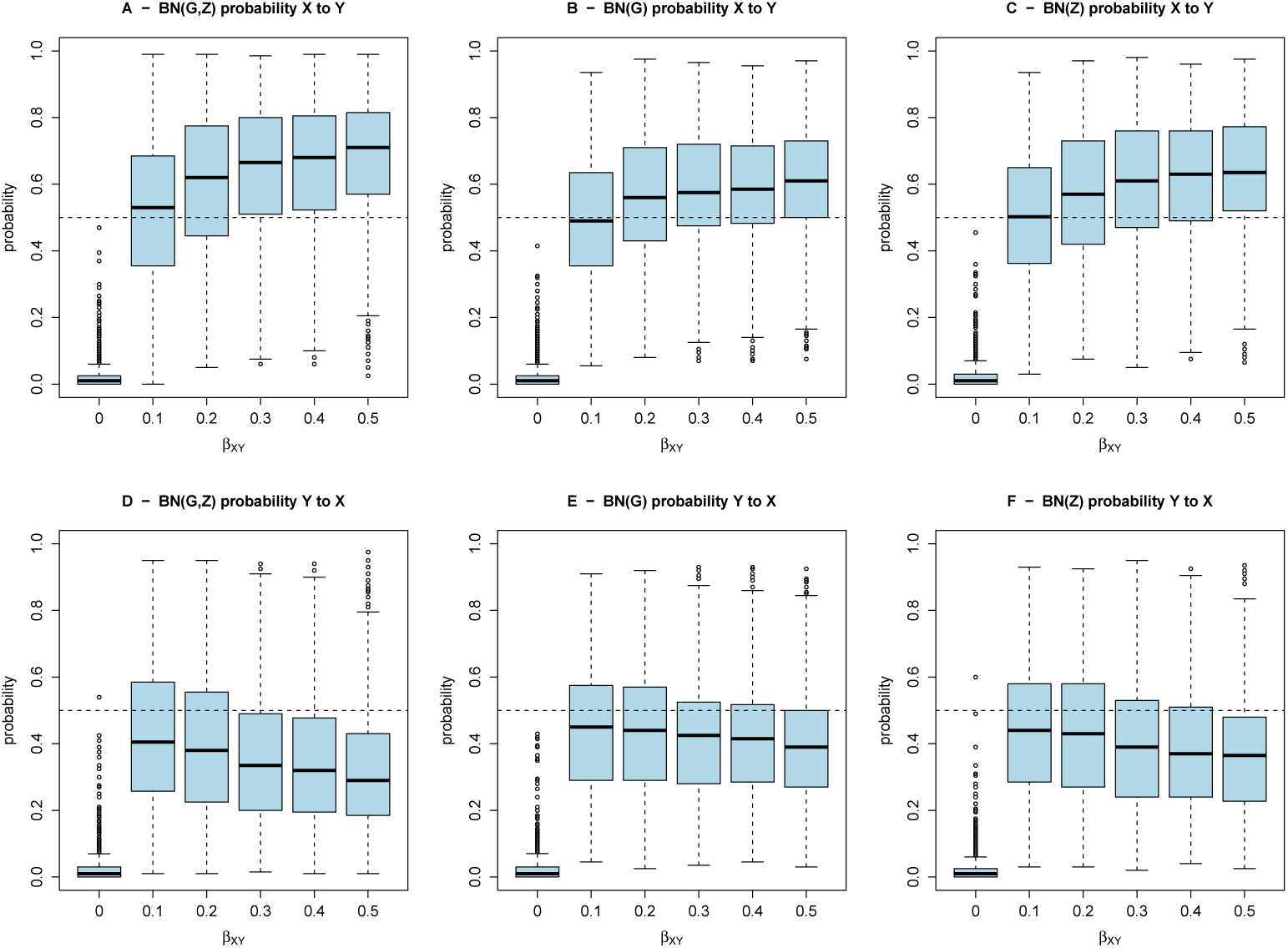
Box plots of estimated BN arrow probabilities for data simulated under model 1.

**S13 Fig.**
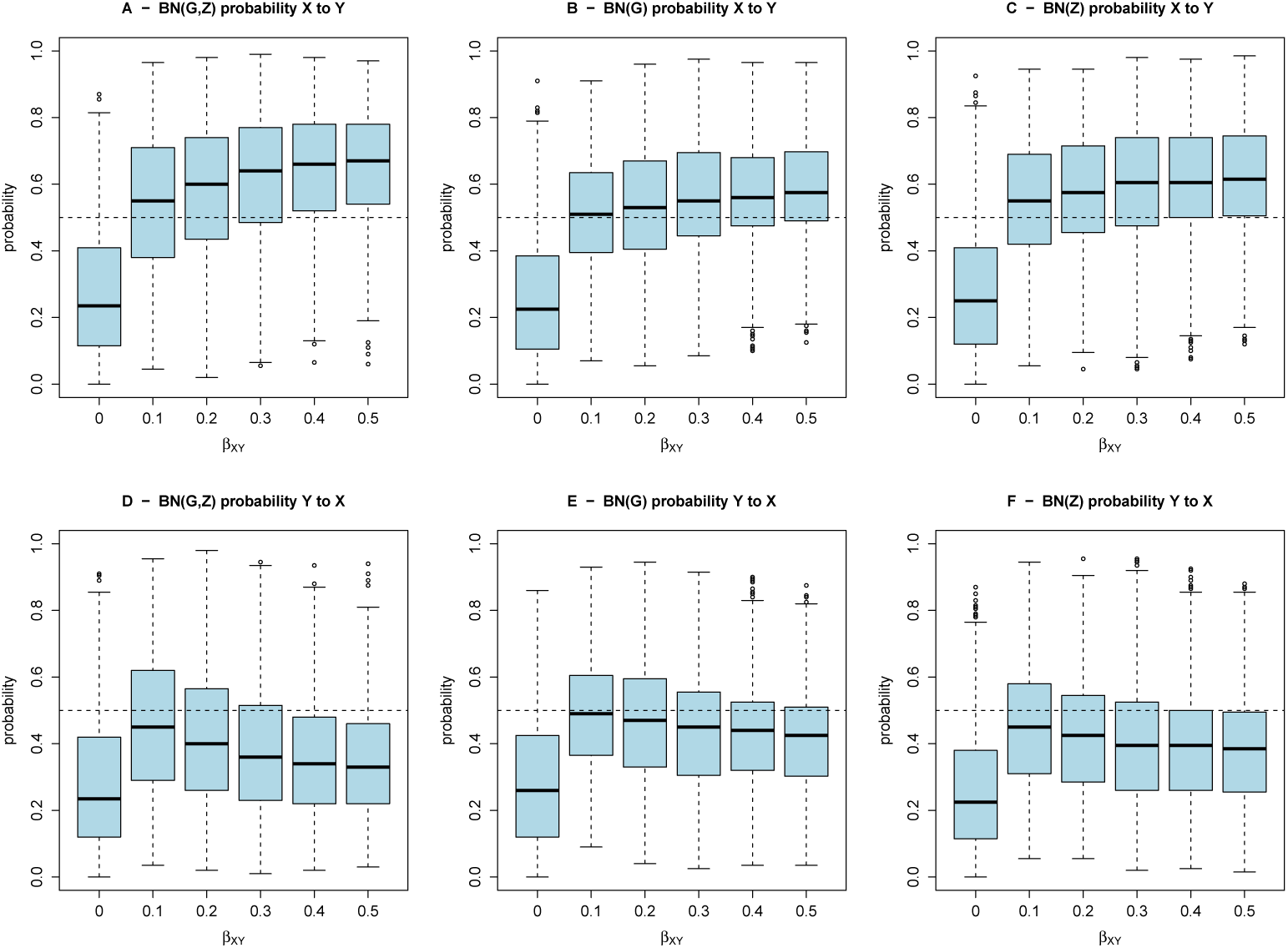
Box plots of estimated BN arrow probabilities for data simulated under model 2.

**S14 Fig.**
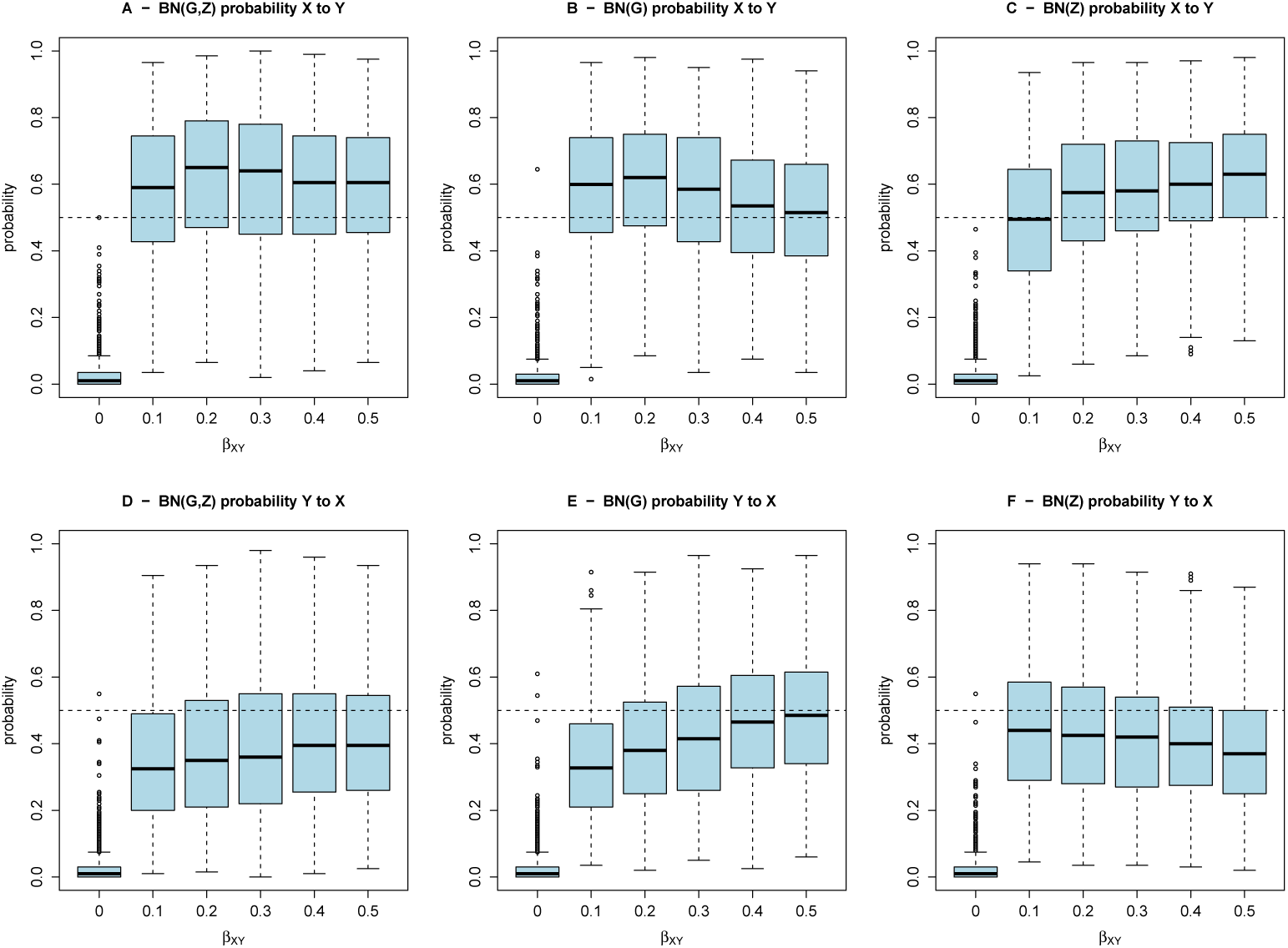
Box plots of estimated BN arrow probabilities for data simulated under model 3.

**S15 Fig.**
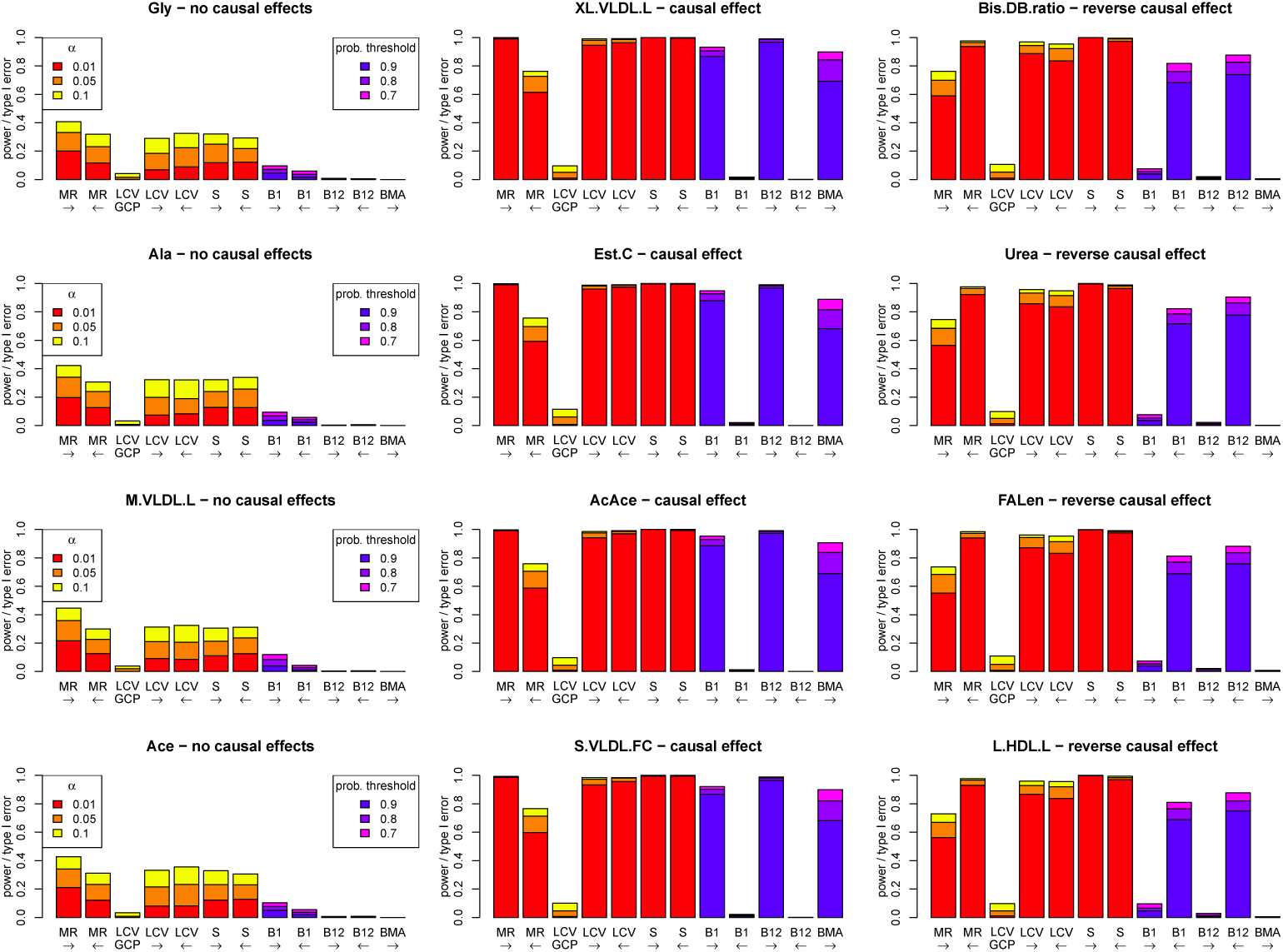
Performance (power and type I error) of different methods under the simulation model used in Figure 10. The model involves 12 metabolites, an outcome Y, 150 SNPs affecting the metabolites, 75 other SNPs affecting Y, and 9775 SNPs with no effect. Four metabolites (middle panels) have a causal effect on Y, four metabolites (right hand panels) have a reverse causal effect from Y to the metabolite, and four metabolites (left hand panels) have no effects to Y in any direction. The left-to-right arrows show tests for a causal effect from the metabolite to Y, and right-to-left arrows show tests from Y to one of the metabolites. MR: Mendelian randomization using an allele score as an instrumental variable for one of the metabolites or Y. LCV: latent causal variable methods where GCP denotes the genetic causality proportion test (testing the null hypothesis that GCP= 0), while LCV with left-to-right or right-to-left arrows corresponds to testing the null hypothesis that GPC= 1 (implying that the metabolite causes trait Y) or that GPC= −1 (implying that Y causes the metabolite), respectively. S: SMUT, using SNPs as random effect variables for one of the metabolites or Y. B1: Bayesian network consisting of one metabolite, Y and the two corresponding allele score variables. B12: Bayesian network consisting of all 12 metabolites, Y and all corresponding allele score variables. BMA: multivariable MR based on Bayesian model averaging (MR-BMA)

**S1 Spreadsheet.** Quantification of data shown in Fig 3.

**S2 Spreadsheet.** Quantification of data shown in Fig 4.

**S3 Spreadsheet.** Quantification of data shown in Fig 5.

**S4 Spreadsheet.** Quantification of data shown in Fig 6.

**S5 Spreadsheet.** Quantification of data shown in Fig 10.

**S6 Spreadsheet.** Quantification of data shown in S1 Fig.

**S7 Spreadsheet.** Quantification of data shown in S2 Fig.

**S8 Spreadsheet.** Quantification of data shown in S3 Fig.

**S9 Spreadsheet.** Quantification of data shown in S4 Fig.

**S10 Spreadsheet.** Quantification of data shown in S5 Fig.

**S11 Spreadsheet.** Quantification of data shown in S6 Fig.

**S12 Spreadsheet.** Quantification of data shown in S7 Fig.

**S13 Spreadsheet.** Quantification of data shown in S8 Fig.

**S14 Spreadsheet.** Quantification of data shown in S9 Fig.

**S15 Spreadsheet.** Quantification of data shown in S10 Fig.

**S16 Spreadsheet.** Quantification of data shown in S11 Fig.

**S17 Spreadsheet.** Quantification of data shown in S12 Fig.

**S18 Spreadsheet.** Quantification of data shown in S13 Fig.

**S19 Spreadsheet.** Quantification of data shown in S14 Fig.

**S20 Spreadsheet.** Quantification of data shown in S15 Fig.

